# Atlas of HIV cis-regulatory elements reveals extensive transcriptional variation across clades, isolates, and within individuals

**DOI:** 10.64898/2026.04.03.716403

**Authors:** Berkay Engin, Mohamed Yousry ElSadec, Joseph Alexander Finkelberg, Tommy Henry Taslim, Daniel L. Bryant, Luis Soto-Ugaldi, Susan Kales, Ching-Huang Ho, Maryam Dashtiahangar, George Muñoz-Esquivel, Elvis Morara, Jacob Purinton, Benedetta D’Elia, Rodrigo Castro, Harshpreet Chandok, Matias Alejandro Paz, Trevor Siggers, John P. Ray, Andrew J. Henderson, Ryan Tewhey, Juan I. Fuxman Bass

## Abstract

Human immunodeficiency virus (HIV) replication, persistence, and reactivation depend on transcription from integrated proviruses. Despite extensive sequence variation, how viral genetic diversity influences transcriptional regulation remains poorly understood. Here, we generate a functional regulatory atlas of HIV-1 and HIV-2 by combining tiling and saturation mutagenesis massively parallel reporter assays (MPRAs) with comparative sequence analysis and predictive modeling. By profiling thousands of HIV isolates in Jurkat and human primary CD4+T cells, we reveal extensive variation in baseline and stimulus-induced long terminal repeat (LTR) activity across and within clades, driven by distinct transcription factor configurations. These activities frequently differ among proviruses from the same individual and shift over infection and transmission without consistent selection for activity. Beyond the LTR, we identify conserved intragenic cis-regulatory elements, revealing regulatory architectures that complement LTR activity. Finally, we develop sequence-based models that accurately predict transcriptional activity, enabling scalable functional annotation of viral diversity and evolution.

## Introduction

Human immunodeficiency virus (HIV), which if untreated can progress to acquired immunodeficiency syndrome (AIDS), establishes persistent infection by integrating into the host genome, generating long-lived proviral reservoirs that sustain infection despite immune clearance or therapy.^1,2,3,4^ Proviral transcription is driven by the long terminal repeat (LTR), whose cis-regulatory architecture enables responsiveness to both activating and repressive cellular signals.^5^ Through these inputs, LTRs toggle between transcriptional activation—promoting viral gene expression and replication—and transcriptional silencing, allowing infected cells to evade immune detection. For instance, the HIV-1 LTR integrates inflammatory and T cell activation signals through transcription factors (TFs) such as NF-κB and AP-1, promoting rapid amplification upon cell activation.^5,6,7,8^

HIV transcription has been investigated for decades, particularly in HIV-1, generating substantial mechanistic insight on LTR activation and silencing. HIV exhibits remarkable genetic diversity: HIV-1 and HIV-2 comprise distinct evolutionary lineages, and HIV-1 group M alone encompasses multiple clades (A–K) and numerous recombinant forms.^9,10,11^ However, most mechanistic studies have relied on a small number of laboratory-adapted or selected strains, resulting in a poor characterization of transcriptional regulatory variation across natural viral diversity. Sequence variation also arises within infected individuals through error-prone reverse transcription and host-driven mutagenesis.^12,13^ The functional consequences of this diversity for LTR transcription—baseline activity, stimulus responsiveness, and the propensity to enter or exit latency—remain difficult to predict and have not been quantified at scale.

Latency establishment and reactivation depend on the interplay between proviral cis-regulatory sequences and host TF environments, and differences among isolates have been proposed to contribute to variation in disease course, transmission, pathogenesis, and reservoir dynamics.^3,14,15^ Cure strategies that aim to reduce the reservoir by inducing proviral expression implicitly assume that genetically diverse proviruses respond comparably. However, differences in TF binding site composition, motif arrangement, and signaling responsiveness across clades and within-host variants may lead to variable reactivation across genetically diverse viral reservoirs, contributing to the limited success of these shock-and-kill strategies.^16,17^

Another major gap is the extent to which HIV transcription is influenced by cis-regulatory elements (CREs) beyond the LTR, embedded within viral coding regions. Intragenic CREs have been described in HIV-1, including elements within *pol* that influence transcription and an *env*-associated promoter capable of initiating transcription independent of the 5′ LTR and contributing to expression from defective proviruses in individuals on antiretroviral therapies (ART).^18,19,20,21,22^ Whether such intragenic CREs are widespread, conserved, or functionally divergent across isolates remains largely unknown.

A systematic approach is needed to quantify transcriptional activity across large numbers of viral sequences, resolve nucleotide-level determinants of activity, and compare regulatory logic across viruses and host contexts. Massively parallel reporter assays (MPRAs) constitute a high-throughput and sensitive approach that measures the transcriptional activity of tens of thousands of sequences in parallel.^23,24,25^ MPRAs have been extensively used to study promoter and enhancer activity of eukaryotic, prokaryotic, and synthetic DNA elements, to identify TF motifs contributing to activity, and to evaluate the effect of genetic variants on gene expression.^23,26,24,25^ More recently, we have used MPRAs to identify and characterize CREs in double stranded DNA viruses, illustrating the broad applicability of this approach.^27^

Here, we define cis-regulatory activity across the genomes of HIV-1 and HIV-2 by integrating tiling and base-resolution saturation mutagenesis MPRAs in Jurkat and human primary CD4+ T cells with large-scale comparative sequence analysis. We systematically quantify how baseline and stimulus-induced transcription varies across thousands of HIV isolates, and define how TF motif number, strength, and spacing impact transcriptional outputs across clades, isolates, and within people with HIV (PWH). We further identify and characterize conserved intragenic CREs in HIV-1 and HIV-2, revealing an additional layer of regulatory control that may complement or compensate for LTR architecture. Finally, we develop sequence-based models that generalize our MPRA data, enabling *in silico* prediction of baseline and inducible LTR activity directly from sequence. This accessible framework supports high-throughput, functional characterization of viral isolates and enables comparative analyses across individuals, transmission pairs, and longitudinal viral evolution. Our experimental data and predictive models are provided as a user-friendly web resource HIVRegDB (https://hivregdb.com/). Together, our studies provide a mechanistic and predictive foundation for understanding how HIV sequence diversity shapes transcriptional behavior across viruses and cellular contexts, with implications for latency modeling and intervention strategies.

## Results

### MPRAs capture key TFs regulating HIV-1 LTR activity

To determine whether MPRAs can be employed to study HIV transcriptional regulation, identify key TFs, and capture activity variation across isolates, we performed three sets of experiments: 1) we tiled the genomes of four HIV isolates (HIV-1: clade B REJO and CH058; and HIV-2: ROD, and GH-1) using overlapping 200 bp tiles offset by 50 bp tested in both orientations, 2) we performed saturation mutagenesis in active regions to identify single nucleotide variants that perturb activity and the TFs involved, and 3) we evaluated activity variation across thousands of isolates from different clades (**Figure 1A** and **Tables S1-4**). MPRA experiments were performed in quintuplicate in Jurkat cells (unstimulated, and treated with 1 µg/ml αCD3 + 10 ng/ml PMA, 20 ng/ml TNFα, or 20 ng/ml IFNγ) and for four human primary CD4+ T cell donors and met stringent quality-control criteria, including barcode complexity, read depth, replicate correlation, and expected activity distributions of positive and negative control sequences (**Figure S1A-G**). MPRA activity in CD4+ T cells strongly correlated with activity measured in unstimulated and αCD3 + PMA–stimulated Jurkat cells (r = 0.88 and 0.92, respectively; **Figure S1H**), showing that Jurkat cells recapitulate the variability in regulatory activity observed in primary cells while providing a scalable system for high-throughput studies.

**Figure 1:**
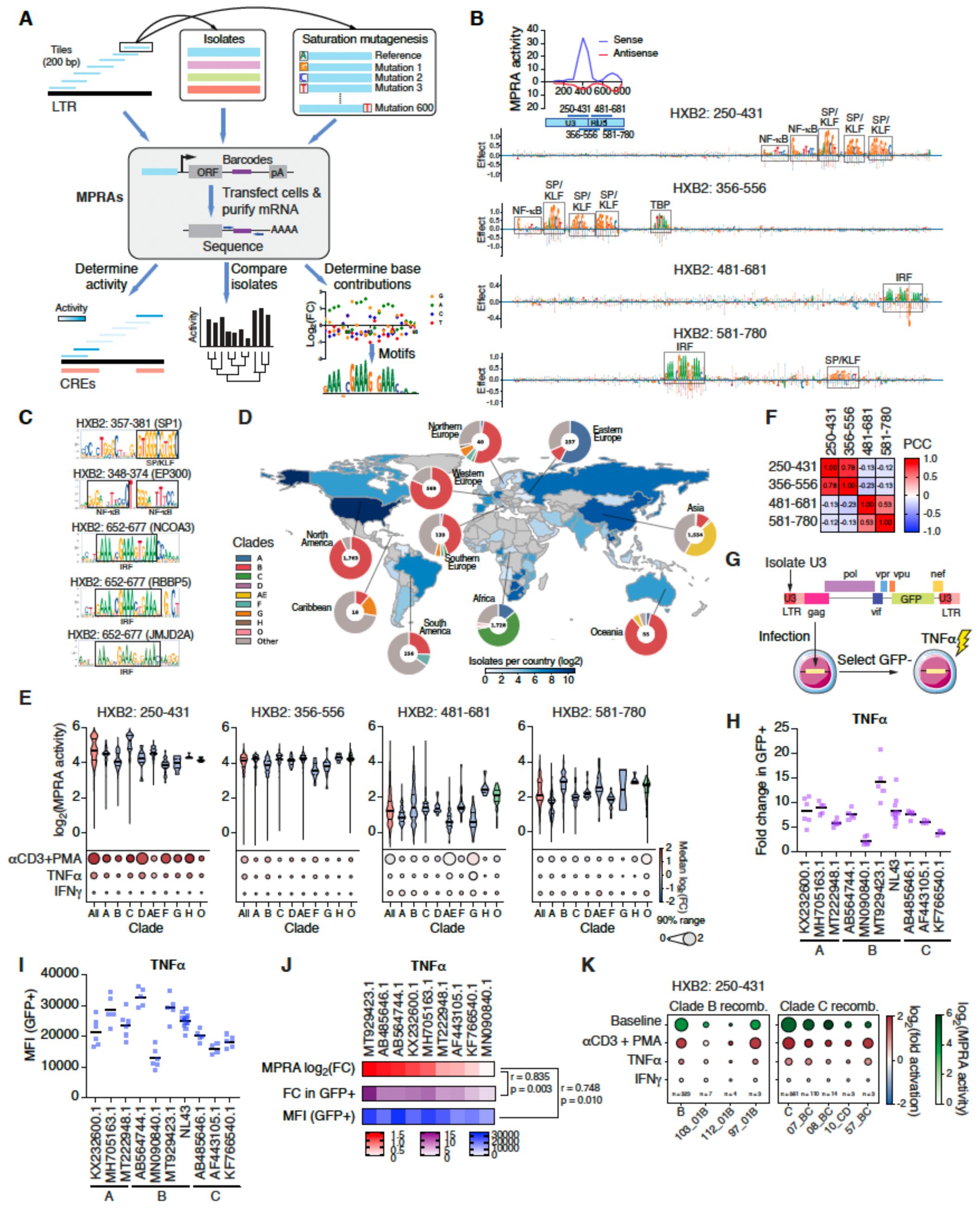
Functional variation of HIV-1 LTRs across clades and isolates. (A) Schematic of MPRAs tiling through HIV genomes, saturation mutagenesis, and evaluation of different isolates. (B) Saturation mutagenesis MPRA experiments in unstimulated Jurkat cells for four tiles in the HIV-1 LTR from the clade B REJO strain that show transcriptional activity. Region coordinates are provided using standardized HXB2 genomic coordinates. TF motifs that contribute to activity are outlined. Inset shows MPRA activity tiling through the HIV-1 LTR. Blue indicates activity of tiles in the sense, and red in the antisense (opposite orientation) strand. (C) CASCADE-derived motifs for different cofactors at various regions of the HIV-1 LTR. (D) Regional distribution of HIV-1 isolates tested by MPRAs. (E) Violin plots showing the distribution of activity in unstimulated Jurkat cells across 5,569 isolates for different regions of the HIV-1 LTR across clades. The thick black line indicates the median, and the dotted lines indicate the first and third quartiles. The dots heatmaps below indicate the fold activation by stimulation of Jurkat cells with αCD3+PMA, TNFα, or IFNγ. The color indicates the median fold activation across isolates, whereas the size of the dots reflects the fold activity differences between the 5th and 95th percentiles. (F) Pearson correlation between the activity of different regions of the HIV-1 LTR across isolates. (G) Schematic of chimeric proviral constructs and reactivation experimental approach. (H-I) Fold change in GFP+ cells relative to DMSO control (H) or geometric mean fluorescence intensity of GFP+ cells (I) in Jurkat cells infected with chimeric proviruses carrying the U3 region of the indicated isolates from clades A, B, and C, and stimulated with TNFα for 2 days. (J) Comparison between fold activation of MPRA data from tile HXB2:250-431 in Jurkat cells activated with TNFα, and provirus reactivation by TNFα measured as fold change in GFP+ cells relative to DMSO control or geometric mean fluorescence intensity of GFP+ cells. Pearson correlation coefficients and one-tailed p-values are indicated for each comparison to MPRA data. (K) Dot heatmaps of recombinant HIV-1 viruses derived from clades B and C. Median baseline activity in tile HXB2:250-431 is shown in shades of green, whereas the median fold activation by αCD3+PMA, TNFα, or IFNγ is shown blue-white-red gradient. The size for the dots reflects the fold activity differences between the 5th and 95th percentiles.

Given the central role of the LTR in HIV transcriptional regulation, we first evaluated whether MPRAs recapitulated known mechanisms underlying LTR activity. HIV LTRs comprise three main regions: (1) U3, containing the modulatory, enhancer, and core promoter domains; (2) R, which includes the transactivation-response RNA element (TAR); and (3) U5, forming part of the leader sequence^5^. Consistent with reports of TF binding site locations,^7^ the HIV-1 U3 enhancer and core-promoter tiles, as well as tiles within U5, displayed high transcriptional activity in Jurkat cells (**Figure 1B**).

Although key TF motifs within the HIV-1 LTR have been identified,^7^ the contribution of individual nucleotides across the element remains incompletely defined. To resolve the regulatory architecture of the LTR at base resolution, we performed saturation mutagenesis MPRA across four active tiles from the HIV-1 REJO isolate, substituting each nucleotide with the alternative three bases (**Figure 1A** and **Table S2**). Activity changes relative to the reference sequence were used to generate base-resolution contribution maps, which were compared with known TF motifs using TomTom.^26,28^ Genomic coordinates are provided as standardized HIV-1 HXB2 coordinates to facilitate comparisons with previous studies and between isolates. This analysis recovered several well-established regulators of LTR activity, including two NF-κB sites, three SP/KLF sites, and the TATA box within U3 (tiles HXB2:250-431 and HXB2:356-556), as well as IRF and SP/KLF motifs in U5 (tiles HXB2:481-681 and HXB2:681-780) (**Figure 1B**).^5,7^ Unlike prior studies that targeted individual binding sites, saturation mutagenesis MPRA quantified the contribution of every nucleotide across active LTR regions, revealing canonical motifs and identifying the G bases within U3 SP/KLF sites as the most mutationally sensitive.

TFs binding to the LTR recruit distinct chromatin-modifying cofactors that modulate proviral transcription. We confirmed several of these TF-cofactor functional interactions using CASCADE, a protein-binding microarray assay that detects cofactor recruitment to DNA elements.^29^ Saturation mutagenesis CASCADE identified motifs corresponding to most TFs detected by MPRAs and revealed recruitment of histone acetyltransferases (p300, NCOA3) and the H3K4 methyltransferase RBBP5, and the histone demethylase JMJD2A, cofactors previously implicated in HIV-1 LTR regulation (**Figure 1C**).^30,31,32,33^ These findings demonstrate that MPRA-identified TF motifs are sufficient to recruit chromatin-modifying cofactors, linking sequence-level regulatory architecture to chromatin-based control of proviral expression. Together, these results establish MPRAs as a robust platform for systematic dissection of HIV-1 regulatory mechanisms.

### Variability in HIV-1 LTR activity across clades and isolates

LTR sequences are highly variable across HIV-1 clades and isolates, reflecting the accumulation of insertions, deletions, and substitutions during viral replication.^12,13^ We hypothesized that such sequence differences influence LTR transcriptional activity and impact latency establishment and reactivation in response to pro-inflammatory cytokines or latency reversal agents. Previous studies based on a limited number of isolates have reported differences in LTR regulation across clades.^14,15,34,35,36,37^ For instance, clade C HIV-1, which accounts for half of global HIV-1 infections and predominates in southern and eastern Africa and India, are considered more active than clade B, prevalent in the Americas, Europe, and Australia, due in part to an additional NF-κB site in U3.^36,37^ To evaluate whether these clade-level differences are broadly observed across natural viral diversity, we analyzed 6,676 HIV-1 isolates with complete genome sequences available in NCBI (**Figure 1D** and **Table S3**). For each isolate, we generated 180–200 bp tiles homologous to the reference tiles in the clade B HIV-1 REJO isolate and measured their activity using MPRAs in Jurkat and human primary CD4+ T cells (**Figure 1A** and **Table S4**). Across isolates, the activity of all four LTR tiles varied substantially, with the central 90% of values spanning a 3.2-8.2 and 5.19-18.9 fold range in Jurkat and CD4+ T cells, respectively (**Figures 1E** and **S2A**).

Consistent with previous reports, our large-scale analysis confirmed that clade C viruses generally displayed the highest activity in U3 (HXB2:250-431) in Jurkat and CD4+ T cells (**Figures 1E** and **S2A**), potentially contributing to the elevated viral loads and transmissibility associated with this clade. We also detected substantial intra- and inter-clade variability in U5 (HXB2:481-681 and HXB2:581-780), with clades B, G, and H exhibiting the strongest activity (**Figures 1E** and **S2A**). Notably, we identified a significant negative correlation between activity in U3 (HXB2:250-431 and HXB2:356-556) and U5 (HXB2:481-681 and HXB2:581-780), suggesting compensatory regulatory mechanisms across LTR regions that may maintain balanced transcriptional output (**Figure 1F**). The measured transcriptional activities across regions and isolates are provided through HIVRegDB, a user-friendly web resource (https://hivregdb.com/).

### Diverse LTR responses to T cell activation across HIV-1 clades and isolates

HIV-1 transcription is highly responsive to extracellular cues, including cytokines and T cell activation, which modulate LTR activity through signal-dependent TFs and directly influence viral replication, reactivation, and disease progression.^2,37^ To determine whether LTRs from different isolates exhibit differential responsiveness to activation or cytokine signaling, we performed MPRAs in Jurkat cells stimulated with 1 µg/ml αCD3 + 10 ng/ml PMA or 20 ng/ml TNFα. Both stimuli strongly induced the activity of tile HXB2:250-431, which typically contains multiple binding sites for NF-κB, a TF downstream of these pathways (**Figures 1B, 1E**, and **S3A-B**). In addition, U5 tiles (HXB2:481-681 and HXB2:581-780) were activated by 20 ng/ml IFNγ (**Figures 1E** and **S3C**). Because IFNγ signals through STAT1 and induces IRF1, and these tiles contain IRF binding motifs (**Figure 1B**), this responsiveness likely reflects direct regulation by interferon-induced TFs. Consistent with this model, saturation mutagenesis MPRAs confirmed enhanced contributions of NF-κB and IRF motifs following their respective stimuli (**Figures S3D-E**).

We next examined activation differences across HIV-1 clades (**Figures 1E** and **S3A-C**). Clade C isolates, which typically contain three NF-κB sites within U3, displayed greater responsiveness to αCD3 + PMA stimulation compared to other clades. In contrast, clade 01_AE—prevalent in Southeast Asia and generally containing a single NF-κB site—exhibited weaker activation following αCD3 + PMA or TNFα treatment (**Figures 1E** and **S3A-B**), consistent with previous reports of reduced TNFα responsiveness of 01_AE LTRs.^35,38^ Notably, 01_AE isolates were more responsive to IFNγ at HXB2:581-780, further supporting a compensatory regulation mediated by IRF-dependent elements.

To determine whether episomal MPRAs recapitulate differences in transcriptional activities between isolates in the context of integrated proviruses, we generated full-length chimeric proviruses in which the U3 region of the HIV-1 clade B NL43 backbone was replaced with U3 sequences from nine different isolates spanning a broad range of activities (**Figure 1G**). These single-round proviruses with eGFP replacing the 5’end of the *env* gene, enable direct monitoring of LTR-driven transcription following integration. Jurkat cells were transduced, GFP-negative cells were sorted two days post-infection and, after 7 days post sorting, cultures were stimulated with TNFα or αCD3 + PMA for 40-48 hours (**Figures S3F-G**). Reactivation was quantified as the fold change in the percent of GFP+ cells relative to DMSO controls and the geometric mean fluorescence intensity of GFP+ cells. Across isolates, we observed marked variation for both frequency and magnitude of expression, even for those belonging to the same clade (**Figures 1H-I** and **S3H-I**). Importantly, fold activation of tile HXB2:250-431 by TNFα and αCD3 + PMA measured by MPRAs correlated significantly with both reactivation frequency and GFP intensity (**Figures 1J** and **S3J**). Together, these results demonstrate that MPRA-based measurements accurately capture isolate-specific differences in stimulus-dependent LTR reactivation potential in the context of integrated proviruses.

### LTRs from recombinant HIV-1 viruses can functionally differ from their parental clades

Co-infection with distinct HIV-1 subtypes can generate recombinant viruses through template switching during reverse transcription, resulting in mosaic genomes that contribute to viral diversity, immune evasion, drug resistance, and global spread.^11,39,40,41^ Whether such recombinants retain the transcriptional regulatory properties of their parental clades remains largely unexplored. We found that HXB2:250-431 regions from recombinant viruses derived from the same parental clade did not necessarily exhibit similar Jurkat baseline activity or responsiveness to αCD3 + PMA or TNFα stimulation (**Figure 1K** and **Tables S4-5**). Notably, LTRs from clade B recombinants displayed substantially greater variability—both among themselves and relative to their parental clade—than those from clade C recombinants. Moreover, although each recombinant class likely originated from a single recombination event, we observed marked activity differences among isolates within several recombinant classes (**Figure 1K**). These findings suggest that LTR activity of recombinants cannot be reliably inferred from parental clade identity and instead exhibit substantial functional diversification.

### Differences in HIV-1 LTR responses are driven by differences in TF configurations

Despite the observed clade-level trends, LTR activity and stimulus responsiveness varied markedly among isolates within the same clade (**Figures 1E** and **S3A-C**). This heterogeneity is expected because HIV-1 clades are defined based on envelope or full-genome phylogeny rather than LTR sequence or function.^42^ Consistent with this, HIV-1 LTRs display extensive diversity in TF binding site configurations (i.e., number and position) across clades and isolates (**Table S6**). U3 typically harbors 1–4 NF-κB sites and 1–4 SP/KLF sites, whereas U5 commonly contains 1–2 IRF, 0–2 E2F, and 0–1 SP/KLF sites (**Figure 2A**). These TF configurations do not strictly segregate by clade, providing a potential explanation for the functional variability observed among isolates within clades (**Figures 1E** and **S2A**). Consistent with prior reports from smaller isolate sets,^37,43^ we found that fold activation by αCD3 + PMA or TNFα in Jurkat cells increased with NF-κB site number, although activity increases modestly beyond two sites (**Figure 2B**). In addition, at least two SP/KLF sites were required for baseline activity as well as for maximum stimulation (**Figure 2C**). Although this may be partially driven by enhanced SP1 activity upon cellular activation,^44^ it may also reflect TF cooperativity in U3.

**Figure 2:**
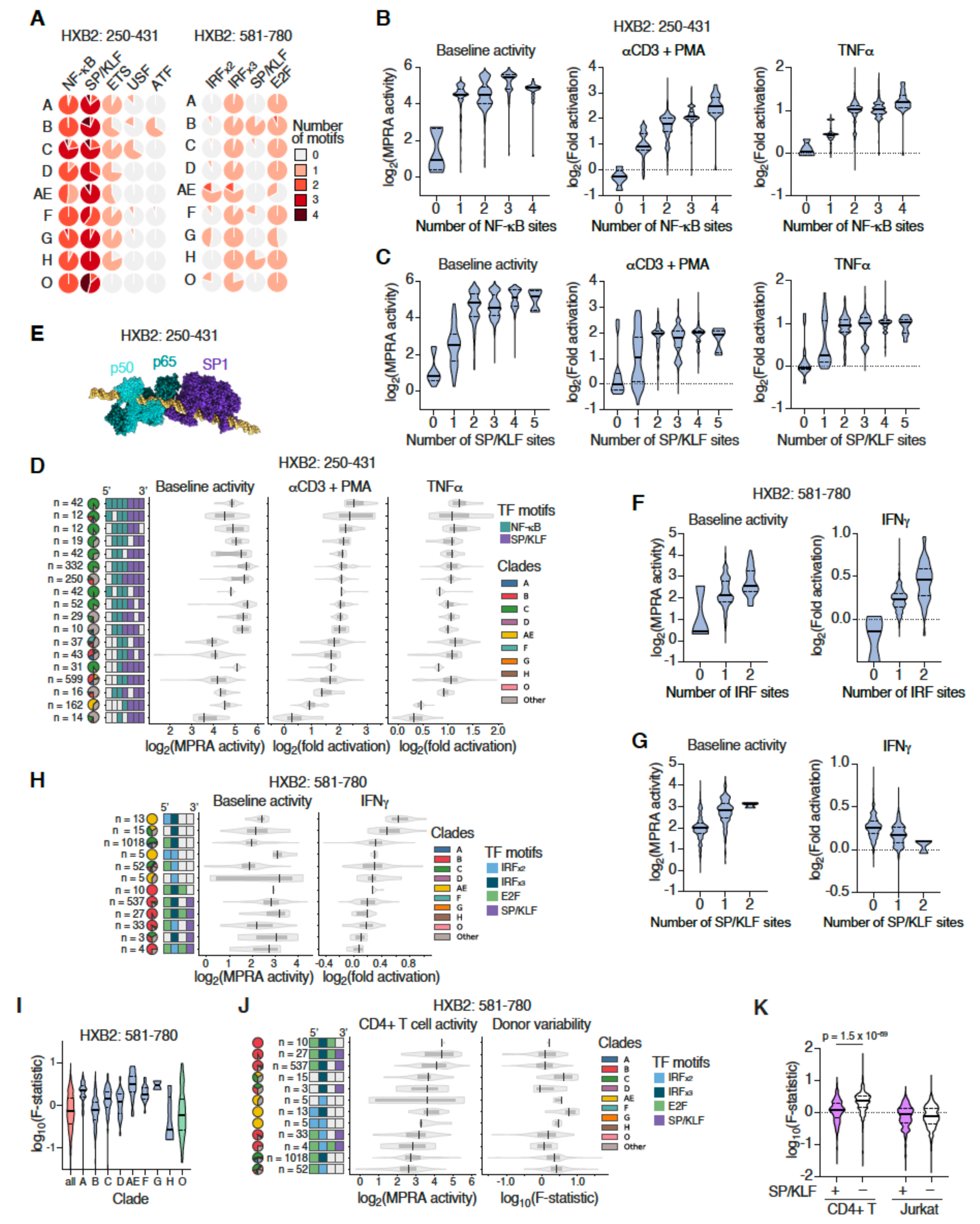
TFs contributing to functional variability between LTRs for different HIV-1 isolates. (A) Distribution of the number of TF binding sites across HIV-1 isolates in tiles HXB2:250-431 and HXB2:581-780. Each pie chart shows the proportion of isolates with different number of binding sites for the indicated TFs. (B-C) Violin plots of baseline activity or fold activation by αCD3+PMA or TNFα for tile HXB2:250-431 across HIV-1 isolates based on the number of NF-κB (B) or SP/KLF sites (C). The thick black line indicates the median, and the dotted lines indicate the first and third quartiles. (D, H) Activity distribution in Jurkat cells across HIV-1 isolates with different TF configurations in tiles HXB2:250-431 (D) and HXB2:581-780 (H). The left boxes represent the TF configurations based on aligned TF positions. The distribution of isolates from different clades across TF configurations is shown as pie chats. Violin plots indicate the distributions of activity in unstimulated Jurkat cells (baseline), and cells stimulated with αCD3+PMA, TNFα, or IFNγ. In the case tile HXB2:250-431 only configurations with at least 10 isolates are shown. (E) Alphafold3 model of the HIV-1 REJO LTR including three SP1 and two sets of NF-κB (p65 and p50) proteins. (F-G) Violin plots of baseline activity or fold activation by IFNγ for tile HXB2:581-780 across HIV-1 isolates based on the number of IRF (F) or SP/KLF sites (G). The thick black line indicates the median, and the dotted lines indicate the first and third quartiles. (I) Violin plots of F-statistic showing the variability in activity for each isolate for tile HXB2:581-780 across four donors in CD4+ T cells. The thick black line indicates the median, and the dotted lines indicate the first and third quartiles. (J) Activity distribution and donor variability (F-statistic) in CD4+ T cells in Jurkat cells across HIV-1 isolates with different TF configurations in tile HXB2:581-780. The left boxes represent the TF configurations based on aligned TF positions. The distribution of isolates from different clades across TF configurations is shown as pie charts. (K) Distribution of donor variability (F-statistic) in tile HXB2:581-780 for isolates that contain or lack an SP/KLF site across four CD4+T cell donors or five replicates of Jurkat cells. Statistical significance determined by two-tailed Brunner-Munzel test.

Given the limited number of isolates assessed in previous studies, the full spectrum of TF configurations within U3 and their impact on LTR activity remained incompletely defined. We therefore mapped NF-κB and SP/KLF site positions within tile HXB2:250-431 across all isolates (**Figure S4A**) and identified configuration-dependent cooperativity. For example, isolates containing NF-κB sites but lacking SP/KLF sites showed little to no inducible activation in MPRAs (**Figures S2B** and **S4B**), consistent with models in which NF-κB activity depends on cooperative interactions with SP1-family factors.^45^ Moreover, among isolates with the same overall site counts, responsiveness differed depending on relative positioning: isolates with one NF-κB and three SP/KLF sites were more strongly induced when the NF-κB site was adjacent to an SP/KLF site (**Figure 2D** and **S4B**). Similar position-dependent effects were observed across additional configurations with matched site number but distinct spacing and arrangement (**Figure 2D** and **S4B**). Structural modeling using AlphaFold3 showed potential protein-protein interactions between SP1 and NF-κB subunits p50 and p65 further supporting the plausibility of physical proximity-dependent NF-κB–SP1 cooperativity at adjacent sites in the REJO LTR (**Figure 2E**). Remaining variability in activity between isolates within NF-κB and SP/KLF configurations likely reflects differences in motif strength and spacing that influence cooperative interactions, as well as the contribution of additional TF motifs within U3.

We also observed TF configuration-dependent differences in Jurkat baseline activity and IFNγ responsiveness within U5 tile HXB2:581-780 (**Figures 2F-H** and **S4C**). IFNγ induction was strongest in isolates containing two IRF sites (**Figure 2F**), and within these, configurations with three (rather than two) GAAA repeats showed greater induction (**Figure 2H**). An SP/KLF site increased baseline activity but reduced IFNγ responsiveness (**Figures 2G-H**). Although adjacent IRF and SP1 sites are known to cooperate at cellular promoters,^46^ the lack of apparent cooperation in tile HXB2:581-780 is consistent with the substantially larger spacing (∼70–90 bp) between these sites in U5 (**Figure 1B**). Together, these results show that stimulus responsiveness across LTRs is strongly shaped by TF configuration and motif arrangement—features that vary extensively across isolates and are not reliably captured by clade designation alone.

### TF configurations influence variability in LTR activity across CD4+ T cell donors

Despite the overall concordance between Jurkat and primary CD4+ T cells, primary cells introduce additional variability driven by donor-specific genetic background, immune history, and cellular state that can affect TF activity and signaling pathways regulating HIV transcription. Inter-donor differences in HIV transcription have been widely reported,^47,48^ but whether such donor-dependent effects influence all viral isolates equally remains unclear. We hypothesized that distinct LTR TF configurations exhibit different sensitivities to donor-specific cellular environments, leading to isolate-specific variability in transcriptional output across donors. To quantify this, we calculated an F-like statistic comparing observed to expected activity variance across donors for each isolate. Whereas tile HXB2:250-431 displayed relatively uniform variability across isolates and clades, tile HXB2:581-780 exhibited substantially higher inter-donor variation (**Figures 2I** and **S2E**). This increased variability was primarily associated with the absence of an SP/KLF site, as isolates containing an SP/KLF motif within HXB2:581–780 showed significantly lower variability across CD4+ T cell donors (**Figure 2J-K**). In contrast, this effect was not observed across Jurkat cell replicates, showing that the reduced variability associated with the SP/KLF motif is not due to technical variation. This is consistent with our observation that SP/KLF sites in U5 dampen IFNγ responsiveness (**Figure 2G**), suggesting that SP/KLF-mediated regulation buffers LTR activity against donor-specific differences in interferon signaling. These results also demonstrate that TF configurations not only shape mean transcriptional output but also influence the robustness of LTR activity across distinct cellular environments.

### Proviruses from the same individuals often vary in LTR activity

Proviral sequences vary substantially within PWH due to mutagenesis by reverse transcriptase, host enzymes such as APOBEC3 and DNA repair factors, insertions and deletions arising during replication, and immune selection.^12,13,49^ This genetic diversity can profoundly affect LTR activity, ranging from transcriptionally inactive defective proviruses to variants with minimal impact on regulatory function.^50,51,52,53^ While large deletions and hypermutation often have predictable consequences, the transcriptional impact of more subtle sequence changes remains difficult to anticipate.

To systematically assess within-host transcriptional variability, we compared baseline activity of tile HXB2:250-431 and its responsiveness to αCD3 + PMA or TNFα stimulation across isolates from 46 PWH obtained from NCBI and Los Alamos Database, each represented by at least five proviral sequences (**Table S3**). Among these individuals, 13 exhibited more than twofold variation in Jurkat baseline or CD4+ T cells activity, including five with defective proviruses lacking detectable transcription (**Figure 3A**). These differences were associated with alterations in SP/KLF sites in 5 of 13 cases, NF-κB sites in 3 of 13 cases, and concurrent changes in both motifs in 5 of 13 cases. Similarly, 18 individuals showed more than twofold variation in inducible activation by αCD3 + PMA or TNFα (**Figure 3A**). Although reduced inducibility was linked to NF-κB site mutations in about half the cases, 22% (e.g., 105_116107) were explained by changes in SP/KLF sites, consistent with cooperativity between NF-κB and SP/KLF.

**Figure 3:**
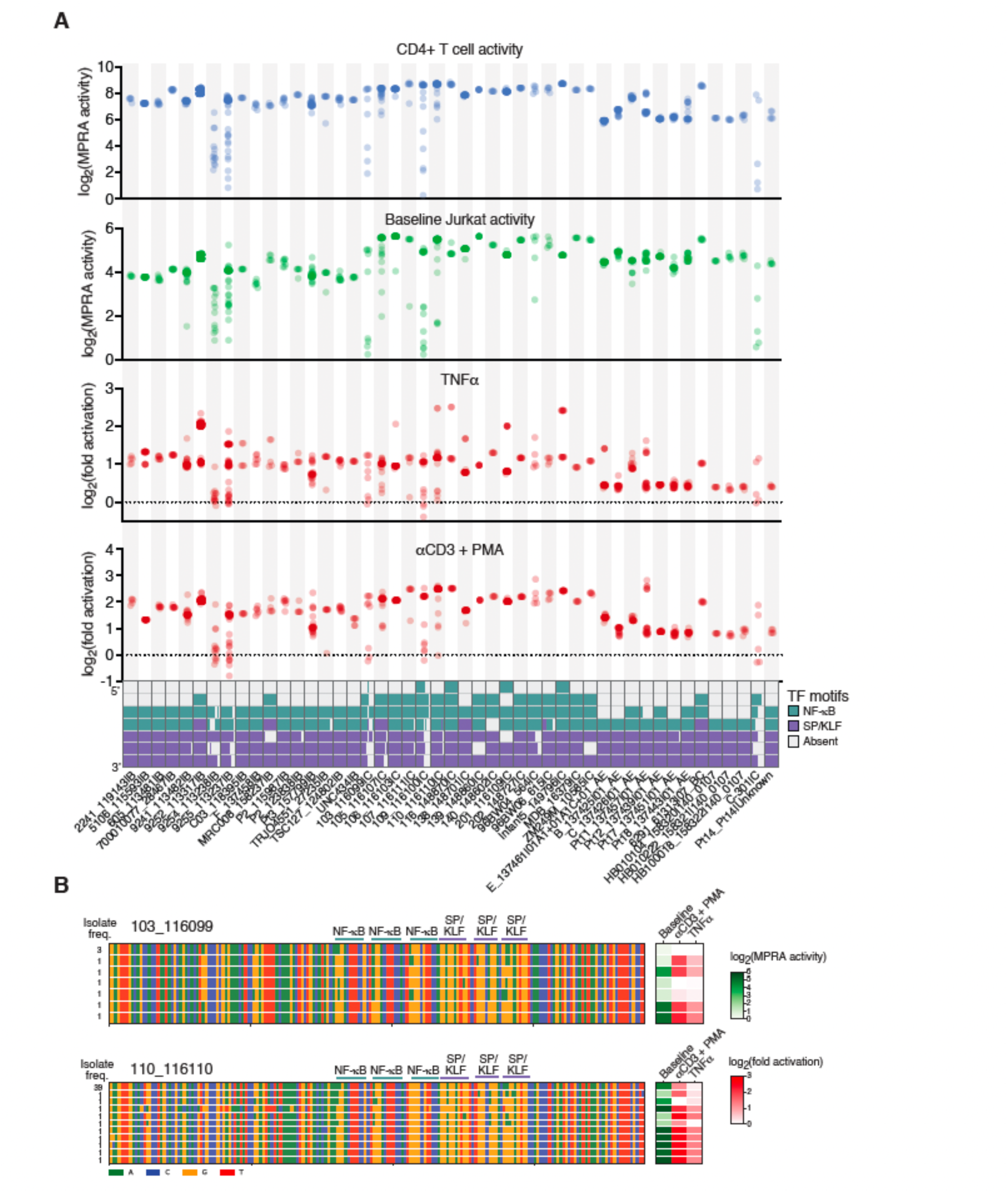
Variation in HIV-1 LTR activity and regulation within PWH. (A) Distribution of baseline activity and fold activation by αCD3+PMA or TNFα for different PWH with at least 5 different isolates in HXB2:250-431. PWH are ordered by HIV-1 clade. The bottom boxes represent the TF configurations based on aligned TF positions across all isolates from the corresponding PWH. (B) Examples of activity and sequence variation for isolates of two PWH from clade C corresponding to tile HXB2:250- 431. The left heatmaps show the sequence alignments for isolates of a PWH. The frequency of isolates with a specific sequence is shown. The binding sites for NF-κB and SP/KLF are indicated. The right heatmap indicates the baseline activity (green) and fold activation by αCD3+PMA or TNFα (red).

G→A hypermutation, a footprint of APOBEC3-mediated cytidine deamination that restricts HIV replication,^13,50^ commonly disrupted GC-rich SP/KLF and NF-κB sites and were associated with both disruption of baseline activity and diminished responsiveness to αCD3 + PMA and TNFα in Jurkat cells (**Figure 3B**). However, this was not universal: proviruses containing sporadic mutations or small deletions sometimes retained high baseline but showed weak inducibility, or vice versa (**Figure 3B** and **S5A**). Mutations occurred across distinct SP/KLF and NF-κB sites and varied extensively both within and between individuals, illustrating how diverse regulatory configurations can arise among proviruses within a single host. We also observed variability in U5 activity among isolates within PWH, driven by mutations affecting IRF or SP/KLF sites (**Figures S5B-C**). Collectively, these findings show that even defective or hypermutated proviruses within the same individual may retain regulatory activity depending on the specific patterns of motif disruption. This functional heterogeneity likely contributes to differences in reservoir persistence and reactivation potential both within and across PWH.

### Development of models to predict activity from sequence

While MPRAs enable detailed assessment of LTR functional diversity, their cost and experimental scale make it challenging to apply them broadly across new samples or on a PWH-specific basis. To address this challenge, we developed two predictive models: 1) CREST (Cis-Regulatory Element Sequence Tester), a deep learning framework that predicts baseline MPRA activity in Jurkat cells from DNA sequence, and 2) LARM (LTR Activation Response Model), a gradient-boosting model that predicts stimulus-induced MPRA fold-activation in response to TNFα for sequences corresponding to tile HXB2:250-431 (**Figure 4A** and **Table S7**).

**Figure 4:**
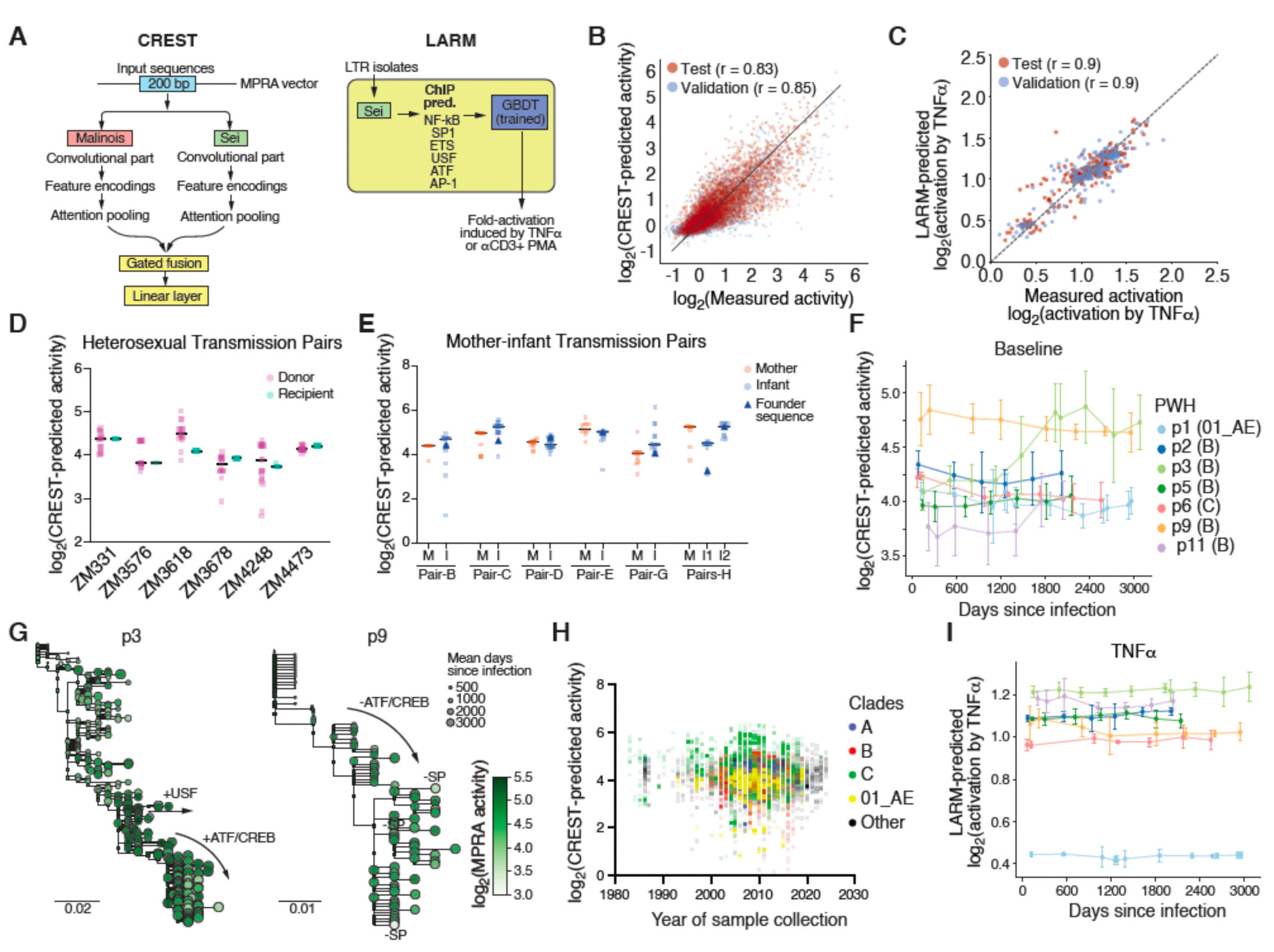
Evolution of HIV-1 LTR activity within PWH using machine learning models. (A) Schematic of CREST and LARM models. CREST predicts baseline activity in Jurkat cells from DNA sequence. LARM predicts fold-activation by αCD3+PMA or TNFα for sequences corresponding to tile HXB2:250-431. (B) Pearson correlation between baseline activity in Jurkat cells measured by MPRAs and predicted using CREST for the validation and test sets. (C) Pearson correlation between TNFα-induced fold activation of tile HXB2:250-431 isolates in Jurkat cells measured by MPRAs and the corresponding values predicted by LARM for the validation and test sets. (D-E) Baseline activity predicted by CREST in tile HXB2:250-431 for isolates heterosexual (D) or mother-to-infant (E) transmission pairs.^62,63^ The activity of the predicted founder sequence is indicated for infants. (F) Baseline activity predicted by CREST in tile HXB2:250-431 for isolates from 7 PWH from samples obtained at different days post infection.^65^ Error bars indicate the standard deviation, dots indicate the average value. (G) Baseline activity predicted by CREST for tile HXB2:250-431 of two PWH. Isolates are shown within their respective evolutionary trees. Node color represents the activity and node size indicates the average day post infection for the corresponding haplotype. Square nodes correspond to predicted ancestral sequences. (H) Baseline activity predicted by CREST for tile HXB2:250-431 for samples from Los Alamos database plotted versus the date of sample collection. Isolates are colored by clade. (I) Fold activation in Jurkat cells stimulated with TNFα predicted by LARM in tile HXB2:250-431 for isolates from 7 PWH from samples obtained at different days post infection. Error bars indicate the standard deviation, dots indicate the average value.

To train CREST, we leveraged an existing MPRA dataset comprising 200 bp genome tiles derived from human double-stranded DNA viruses^27^. Given the relatively modest size of this dataset, we adopted a transfer-learning approach. As a pretrained source model, we used Malinois, a previously developed MPRA predictor trained across three cell types: K562, HepG2, and SK-N-SH.^26^ To capture Jurkat-specific transcriptional regulators whose binding preferences are not learned by Malinois, we incorporated Sei as a second source model,^54^ which predicts chromatin binding profiles for hundreds of proteins across multiple cell types, including Jurkat. In total, 49,316 tiles were used for training, 8,653 for validation, and 7,059 for testing (**Table S7**). All retroviral tiles were excluded from the training set to enable an unbiased assessment of model performance on these viruses.

CREST predictions showed strong agreement with experimentally measured MPRA activity in both validation and test sets (r=0.85 and r=0.83, respectively; **Figure 4B**). High predictive performance was observed across most HIV sequences, with correlations of 0.84-0.92 (**Figure S6A**). CREST also distinguished activity differences among closely related LTR variants, as illustrated by strong correlation between predicted and measured activity for tile HXB2:250-431 across HIV-1 isolates (r=0.72; **Figure S6B**). Moreover, *in silico* saturation mutagenesis using CREST recovered TF motifs identified experimentally by saturation mutagenesis MPRAs (**Figure S6C**), indicating that the model learns biologically meaningful regulatory features. Collectively, these results demonstrate that CREST accurately predicts Jurkat MPRA activity across diverse viral sequences and captures functional motif-level determinants.

Because the probability and magnitude of proviral reactivation correlated with MPRA fold activation for tile HXB2:250-431 (**Figure 1J** and **S3J**), we next sought to develop a model specifically optimized for predicting stimulus-induced responses. Given the limited number of activation-responsive sequences available for training and the high sequence similarity across isolates within this tile, we adopted a simpler, interpretable model architecture. LARM employs a two-step modeling approach. First, Sei^54^ was used to predict ChIP-seq-like tracks for TFs implicated in LTR regulation, including ATF, SP, NF-κB, ETS, and USF. This yielded 278 Sei-derived features representing isoform- and cell-type-specific occupancy preferences. Second, these features were used to train a gradient-boosting decision tree model to predict fold activation, with hyperparameters manually tuned to improve performance.

When trained on MPRA activation data from Jurkat cells stimulated with TNFα or αCD3+PMA, LARM achieved strong predictive accuracy across validation and test HIV-1 isolates (**Figures 4C and S6D** and **Table S7**), capturing isolate-specific differences in inducible activity. More importantly, LARM predictions also significantly correlated with both the fold increase in reactivated GFP-positive cells and the geometric mean fluorescence intensity in Jurkat cells infected with single-round proviruses harboring distinct LTR sequences (**Figures S6E-H**). Thus, LARM accurately predicts transcriptional response in episomal assays and correlates strongly with proviral expression. CREST and LARM predictions and CREST-derived motifs are available in https://hivregdb.com/, which also supports activity prediction for user-submitted sequences.

### CREST predicts limited selection for LTR activity during HIV-1 transmission

HIV-1 transmission is characterized by a strong genetic bottleneck, with most new infections established by a single or limited number of transmitted viruses.^55,56,57^ This bottleneck has been associated with selection for specific viral phenotypes, including CCR5 tropism, envelope properties that enhance mucosal transmission, and resistance to innate immune factors.^56,58,59^ While viral load and replicative capacity are known determinants of HIV transmission,^60,61^ whether variation in LTR-driven transcription also contributes to transmission fitness remains unclear.

Because baseline transcriptional activity may affect viral replication kinetics, antigen expression, and immune clearance, we hypothesized that transmission might favor either stronger LTRs (to enhance early replication) or weaker LTRs (to reduce immune detection during establishment of infection). To test this, we used CREST to predict baseline Jurkat activity of tile HXB2:250-431 across sequences derived from epidemiologically linked heterosexual transmission pairs as well as mother-infant transmission cohorts (**Table S8**).^62,63^ Across heterosexual transmission pairs, recipient isolate activity fell within the range of donor isolates, with no evidence that transmission favors higher- or lower-activity variants within the donor population (**Figure 4D**). Similarly, analysis of mother-infant pairs also showed no systematic enrichment of high- or low-activity LTRs in transmitted/founder viruses (**Figure 4E**). Although the number of pairs may limit power to detect subtle directional effects, these observations do not support a strong role for intrinsic LTR transcriptional strength in determining transmission fitness. In contrast to the lack of bias in LTR activity, prior studies of envelope-mediated phenotypes have identified clear signatures of selection during transmission, consistent with a dominant role for viral entry, immune evasion, and mucosal fitness in shaping the transmission bottleneck.^56,57,58^ Our results also align with studies showing that early transmitted viruses do not consistently display enhanced replication capacity relative to chronic viruses in baseline conditions.^58^

### CREST and LARM predict longitudinal evolution of plasma HIV-1 LTR activity

HIV-1 transcription is tightly linked to viral replication, immune recognition, and pathogenesis, placing the LTR under strong and potentially opposing selective pressures during infection. While reduced transcription can limit viral replication and competitive fitness, it diminishes immune targeting, whereas elevated transcription can increase viral replication but also antigen expression, potentially enhancing immune-mediated clearance.^3,64^ Whether one of these trade-offs predominates, and whether selective pressures act similarly across individuals over the course of infection, has not been systematically examined at the transcriptional regulatory level.

To directly assess longitudinal evolution of LTR regulatory activity, we evaluated HIV-1 genome sequences from biobanked plasma samples collected over time from 7 PWH (five from clade B, one clade C, and one clade 01_AE). All individuals were treatment-naive with well-estimated dates of infection and were diagnosed between 1990 and 2003.^65^ Each individual contributed 6-12 plasma samples spanning early infection [∼33–209 days after estimated date of infection (EDI)] through late follow-up (∼5.5-8.3 years post EDI), yielding 73 total samples and 1,139 LTR sequences that capture actively replicating viral populations. We applied CREST and LARM to predict baseline activity and fold activation by TNFα in U3 (HXB2:250-431) (**Table S9**).

Across individuals, baseline U3 MPRA activity exhibited heterogeneous evolutionary trajectories. Of the seven PWH, three showed consistent decrease in predicted activity over time, two showed consistent increase, and two displayed no significant directional change (**Figures 4F** and **S6I**). For example, late isolates from p3 generally exhibited higher activity than earlier sequences across most branches of the LTR phylogeny, driven by the gain of ATF/CREB and USF binding sites in distinct lineages (**Figure 4G**). In contrast, late isolates from p9 showed progressive reductions in activity over time, associated with loss of ATF/CREB sites in the lineage leading to later timepoints, as well as sporadic losses of an SP/KLF site (**Figure 4G**). While changes in TF binding site composition contribute to this variation, additional mechanisms such as chromatin regulation and transcriptional elongation dynamics may also influence LTR activity. Together, these findings indicate the absence of a uniform selective pressure acting on LTR transcriptional strength across individuals and likely reflect distinct balances between viral replication efficiency and immune-mediated clearance in different hosts which may depend on individual immune histories. This lack of consistent directionality in activity during transmission and longitudinal evolution also helps explain why the average U3 activity has remained relatively stable within clades across the broader HIV-1 pandemic (**Figure 4H**).

We next examined predicted inducible activation of U3 (HXB2:250–431) in response to TNFα using LARM. In contrast to baseline activity, fold activation remained largely stable over time within each individual, with minimal variability across longitudinal samples (**Figures 4I** and **S6J**). Given that LARM predictions correlate strongly with experimentally measured reactivation of integrated proviruses (**Figures S6E-H**), these results suggest that while constitutive LTR activity can evolve over the course of infection, responsiveness to activation cues is comparatively constrained once infection is established. Finally, because our analyses were based on plasma-derived viral RNA, the observed trajectories reflect evolutionary dynamics within actively replicating viral populations and may not capture potential regulatory changes occurring within the latent proviral reservoir.

### Systematic Identification of conserved intragenic CREs in the HIV-1 genome

Previous studies have identified intragenic CREs within the HIV-1 genome with diverse functional roles.^18,19,20,21,22^ For instance, an enhancer within *pol* that recruits Sp1, AP-1, Oct-1/2, and PU.1 promotes proviral transcription.^18,19,20,21^ More recently, we identified a promoter in *env* that initiates transcription independently of the 5′LTR and contributes to expression from defective proviruses.^22^ However, whether additional intragenic CREs are present in the HIV-1 genome and whether such elements are conserved across clades and isolates remains largely unexplored.

To systematically identify intragenic CREs, we performed MPRAs in Jurkat cells using 200 bp tiles with 50 bp offsets across the HIV-1 genome (clade B REJO isolate), testing each tile in both orientations (**Figure 1A** and **Table S1**). Consistent with previous reports, we detected the known CREs within *env* (HXB2:7784-7984) and *pol*, and an additional CRE within *gag* (HXB2:1330-1530) (**Figure 5A**). Saturation mutagenesis MPRAs in Jurkat cells revealed that the CRE in *gag* is regulated by ETS and bZIP motifs, whereas the CRE in *env* is controlled by ETS, CREB/ATF, and ZBED4 motifs (**Figure 5B**). In primary CD4+ T cells, saturation mutagenesis experiments further identified two CEBP motifs within the CRE in *env* (**Figure 5B**). In contrast, no clear TF motifs were detected in the CREs in *pol*, and high-contribution regions overlapped predicted 3′ splice sites, a known RNA processing artifact in plasmid-based reporter systems;^66^ these *pol* regions were therefore excluded from subsequent analyses.

**Figure 5:**
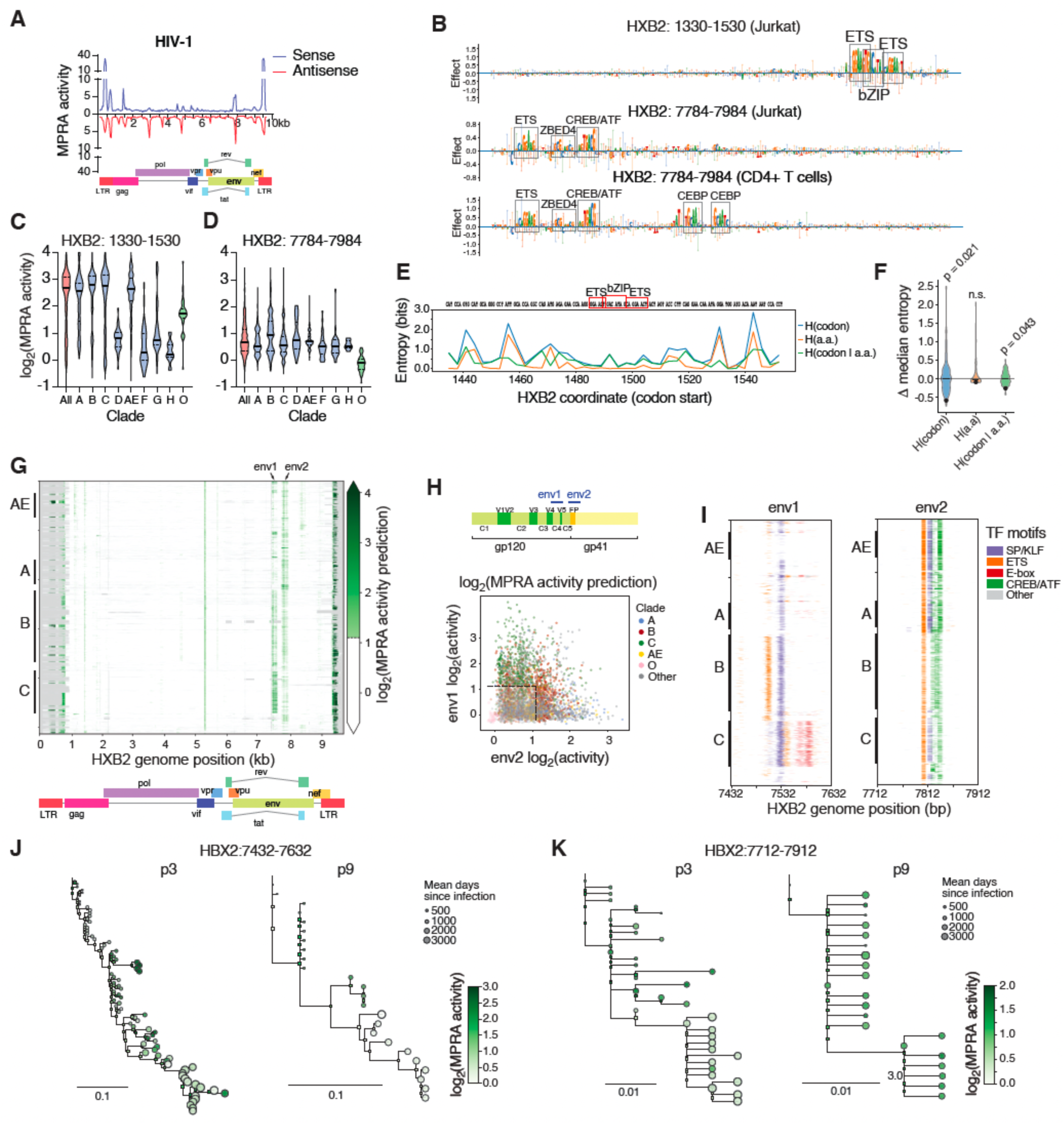
Identification and characterization of intragenic CREs in the HIV-1 genome. (A) MPRA activity map across the genome of HIV-1 clade B REJO strain in unstimulated Jurkat cells. The genome organization is shown below. (B) Saturation mutagenesis MPRA experiments in unstimulated Jurkat cells for HIV-1 intragenic CREs at HXB2:1330-1530 and HXB2:7784-7984. TF motifs that contribute to activity are outlined. (C-D) Violin plots showing the distribution of activity across isolates for intragenic CREs from HIV-1 across clades: HXB2:1330-1530 (C) and HXB2:7784-7984 (D). The thick black line indicates the median, and the dotted lines indicate the first and third quartiles. (E) Entropy decomposition across the ETS–bZIP region in HIV-1 Gag aligned to HXB2 coordinates (codon start positions). Codon entropy (H(codon)), amino-acid entropy (H(a.a.)), and conditional synonymous entropy (H(codon|a.a.)) are shown; the motifs are outlined. (F) Violin plots showing the distribution of Δ median entropy (ROI − rest of window) from circular-shift permutation tests for H(codon), H(a.a.), and H(codon|a.a.). P values are indicated; n.s. denotes not significant. (G) Los Alamos National Laboratory (LANL) genome alignments of HIV-1 strains using the HXB2 genome as a reference. Transcriptional activity predicted using CREST is shown in shades of green. (H) Scatter plot showing the activity predicted using CREST for the two CREs in HIV-1 *env*. Dots are colored by clade. The top diagram shows the location of the CREs within the *env* protein coding sequence. C1-5 = conserved regions, V1-5 = variable regions, FP = fusion peptide. (I) TF motif profiles determined using CREST-based saturation mutagenesis across HIV-1 isolates from LANL. Regions were aligned to the HXB2 genome. (J-K) Baseline activity in Jurkat cells for two PWH predicted using CREST corresponding to HXB2 regions 7432-7632 (J) and 7712-7912 (K). Isolates are shown within their respective evolutionary trees. The color of the nodes represents activity levels. The size of the nodes indicate the day post infection of the corresponding sample. Square nodes represent predicted ancestral sequences.

Given the genetic diversity of HIV-1, we next examined whether the activity of intragenic CREs also varied among isolates. The *gag* CRE was predominantly active in clades A, B, C, and 01_AE, and was absent or weakly active in other clades (**Figures 5D** and **S2D**). In contrast, the *env* CRE generally showed low/moderate activity across clades. The activity of these CREs was also variable within clades and even across isolates from the same PWH (**Figure S7**). This pattern is consistent with intra-PWH variation being enriched among proviral DNA-derived sequences, whereas viral RNA from plasma showed comparatively less variability (**Figures S7E-F**), suggesting that defective proviruses may be a major source of within-PWH regulatory heterogeneity.

To evaluate whether the CRE in *gag* imposed evolutionary constraints beyond amino acid conservation, we quantified entropy in codon usage and in codon usage conditioned on amino acid identity. The region overlapping the ETS and bZIP motifs showed significantly reduced entropy (**Figure 5E-F**), indicating nucleotide-level constraint consistent with purifying selection acting on regulatory function in addition to protein coding requirements. We were unable to perform a comparable analysis for the CRE in env because this region overlaps the antisense open reading frame (asp), introducing additional evolutionary constraints acting on the opposite strand.

Because initial *env* and *pol* CRE discovery was performed using a single clade B isolate, we next applied CREST to full-length HIV-1 genomes from the Los Alamos National Laboratory database to systematically identify intragenic CREs across clades and isolates. CREST recovered the experimentally identified CREs in *gag* and *env* (**Figure 5G**). In addition, we uncovered two additional active regions in *vif* (HXB2:5240-5450) and *env* (HXB2:7432-7632). Saturation mutagenesis of the region in *vif* did not produce known TF motifs, but rather a 3’ splice site, suggesting a likely prediction artifact, and was therefore excluded from subsequent analyses.

The two CREs in *env* exhibited clade-specific activity patterns predicted by CREST: the upstream element (env1) was more frequently active in clade C, whereas the downstream element (env2) was more active in clades A and B (**Figures 5G-H**). Both CREs in *env* were absent in outgroup O viruses, suggesting that they emerged early during the evolution of group M (**Figure 5H**). *In silico* saturation mutagenesis using CREST showed distinct motif architectures across clades and isolates (**Figure 5I**). Whereas env1 was regulated by an SP/KLF site across most isolates, other sites were more variable: clade B env1 is also regulated by an upstream ETS site, and clade C env1 is regulated by downstream ETS and E-box sites. Env2 was predominantly regulated by ETS, CREB/ATF, and SP/KLF (except for Clade B) sites across most isolates (**Figures 5B** and **5I**). Notably, these CREs overlap highly conserved regions of *env*, with env1 residing within the C4 region of gp120 and env2 overlapping the fusion peptide region of gp41, linking regulatory evolution with structurally constrained protein domains.

Longitudinal analysis of activity in the CREs in *env* across phylogenetic branches within individual PWH^65^ revealed dynamic but non-directional changes in regulatory output (**Figures 5J-K**). This lack of directional evolution suggests that regulatory changes within CREs in *env* are not selected for transcriptional output but instead may either be neutral or arise as a byproduct of selective pressures acting on the encoded envelope protein. Given that the *env* region contains transcription start sites independent of the 5′ LTR and that defective proviruses can generate intragenic transcripts associated with immune dysfunction, ^22,67,68^ sporadic CRE activity within *env* may represent an underappreciated source of persistent viral transcription that contributes to HIV-1 pathogenesis despite ART.^67^

### Regulatory similarities and differences between HIV-1 and HIV-2

HIV-2 infects ∼1-2 million individuals, primarily in West Africa, and is characterized by lower transmissibility and slower disease progression compared with HIV-1.^69,70,71,72^ These clinical differences have been associated with reduced replication levels, inefficient entry, and more effective immune control.^69,70,71,72,73^ Previous studies have suggested that HIV-2 LTRs are less transcriptionally active than HIV-1 LTRs, largely due to differences in TF motif configurations, including the presence of a single NF-κB site.^74,75,76^ Tiling MPRA experiments across the HIV-2 ROD strain LTR showed activity in U3 but not in U5, consistent with studies showing that TF motifs in the HIV-2 LTR are restricted to U3 (**Figure 6A** and **Table S1**).^74^

**Figure 6:**
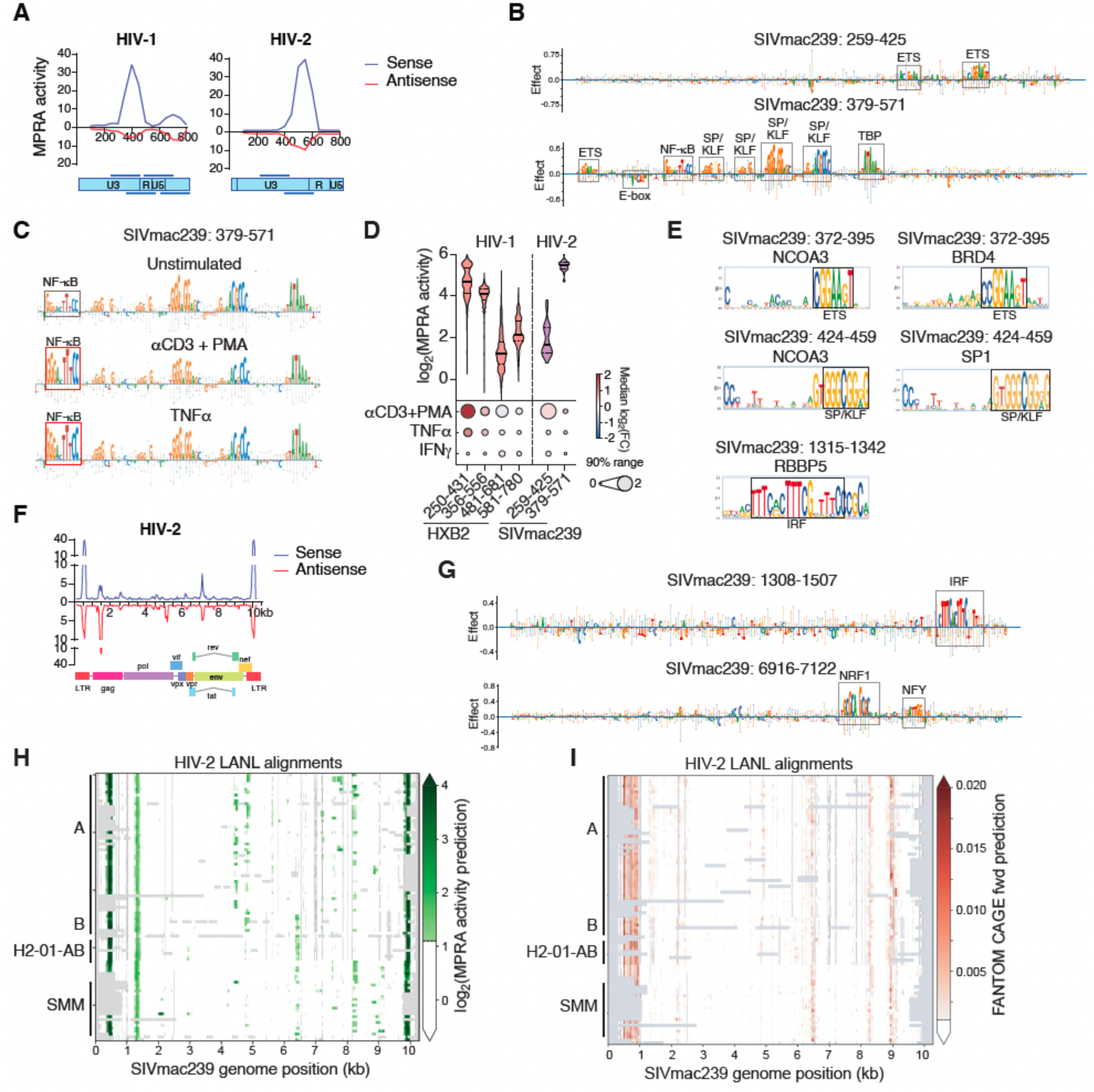
Comparison between HIV-1 and HIV-2 transcriptional regulation. (A) MPRA activity in Jurkat cells tiling through the LTRs of HIV-1 and HIV-2. (B, G) Saturation mutagenesis MPRA experiments in unstimulated Jurkat cells for tiles in the LTR (B), *gag/env* (G) regions of HIV-2 ROD strain that show transcriptional activity. Region coordinates are provided using standardized SIVmac239 genomic coordinates. TF motifs that contribute to activity are outlined. (C) Saturation mutagenesis MPRA experiments in unstimulated and stimulated Jurkat cells for tile SIVmac239:379-571 in the HIV-2 LTR. (D) Violin plots showing the distribution of LTR activity in unstimulated Jurkat cells across 41 HIV-2 isolates with full-length sequences in NCBI. The thick black line indicates the median, and the dotted lines indicate the first and third quartiles. The dots heatmaps below indicate the fold activation by stimulation of Jurkat cells with αCD3+PMA, TNFα, or IFNγ. The color indicates the median fold activation across isolates, whereas the size of the dots reflects the fold activity differences between the 5th and 95th percentiles. (E) CASCADE-derived motifs for different cofactors in the LTR and *gag* regions of HIV-2 ROD strain. (F) MPRA activity map across the genome of HIV-2 ROD strain in unstimulated Jurkat cells. The genome organization is shown below. (H-I) Los Alamos National Laboratory (LANL) genome alignments of HIV-2 strains using the SIVmac239 genome as a reference. (H) Transcriptional activity predicted using CREST is shown in shades of green. (I) Transcription start sites predicted using Puffin trained on FANTOM CAGE data are shown in shades of red.

Saturation mutagenesis MPRAs recovered all known TF sites in the HIV-2 LTR (provided in SIVmac239 coordinates), including one NF-κB site, two ETS sites, and four SP/KLF (one inverted) sites, and the TATA box (**Figure 6B** and **Table S2**).^74,75^ As in HIV-1, the NF-κB site contributed more strongly to activity following αCD3+PMA or TNFα stimulation (**Figure 6C**). However, the magnitude of induction was lower than that observed for most HIV-1 clades containing multiple NF-κB sites and was comparable to clade 01_AE, which also harbors a single NF-κB site (**Figures 6D** and **S3A-B**).

Using CASCADE, we found that SP/KLF sites recruit SP1, similar to HIV-1 (**Figure 6E**). In contrast to HIV-1, where NCOA3 recruitment is mediated through IRF sites in U5, NCOA3 was recruited to the HIV-2 LTR through ETS and SP/KLF sites (**Figure 6E**). ETS sites also recruited BRD4, a known regulator of HIV-1 transcription whose role in HIV-2 regulation remains unclear. In addition to the known regulatory motifs, we identified a repressive E-box upstream of the NF-κB site in the HIV-2 LTR (**Figure 6B**). Similar E-box elements have been found in the HIV-1 LTR; however, these flank the TATA box and can activate or repress transcription, depending on the E-box factors engaged.^77^ Given that repressive E-box TFs such as TCF12 and TFAP4 recruit HDACs and CtBP,^78,79^ the E-box site in HIV-2 may contribute to LTR silencing dynamics and lower overall transcriptional activity. Together, these findings indicate that HIV-1 and HIV-2 share core transcriptional mechanisms but differ in TF usage and configuration.

In contrast to HIV-1, and consistent with the absence of activity in the HIV-2 U5, we did not detect IRF motifs in the HIV-2 LTR (**Figure 6B**). We therefore investigated whether IRF-regulated CREs might exist outside the LTR. MPRA tiling across the full HIV-2 ROD genome revealed two intragenic CREs (**Figure 6F**). One CRE within *gag* (ROD aligned to SIV coordinates: SIVmac239:1308-1507) exhibited high activity, contained an IRF motif, and recruited the H3K4 methyltransferase RBBP5, paralleling IRF-mediated regulation observed in HIV-1 U5 (**Figures 6E** and **6G**). Conversely, whereas ETS regulation occurs in the HIV-2 LTR, the HIV-1 CRE in *gag* is regulated by ETS motifs (**Figure 5B**). These findings suggest that regulatory differences between HIV-1 and HIV-2 LTRs may be partially compensated by intragenic CREs proximal to the LTR.

We also identified an intragenic CRE in *env* (SIVmac239:6916-7122) (**Figure 6E**); however, this region is not homologous to the HIV-1 CREs in *env* and is regulated by distinct TFs (**Figure 5B and 6G**). Whereas the CRE in HIV-1 *env* is controlled by ETS, CREB/ATF and CEBP, the CRE in HIV-2 *env* was regulated by NFY and NRF1, indicating divergent regulatory mechanisms. Genome-wide activity prediction using CREST across HIV-2 isolates from Los Alamos National Laboratory database, revealed strong conservation of the CRE in *gag* but greater variability in the CRE in *env* (**Figure 6H**). This analysis further uncovered additional putative intragenic CREs in *env*, *pol*, and *vpr*, as well as several predicted transcription start sites (**Figure 6I**), highlighting a diverse intragenic regulatory landscape in HIV-2.

## Discussion

HIV transcription has largely been characterized using a small number of reference isolates and individual TF binding sites, yielding deep mechanistic insight but a limited understanding of how regulatory architectures vary across naturally circulating viruses across and within PWH. By combining high-throughput reporter assays with comparative and predictive modeling, our study quantifies transcriptional activity at scale, revealing that HIV regulation is far more diverse and evolutionarily dynamic than previously appreciated. LTRs have flexible regulatory architectures whose transcriptional output depends on TF motif number and arrangement as well as compensation by intragenic regulatory elements. This regulatory diversity shapes baseline transcriptional strength, responsiveness to cellular activation, robustness across host environments, and evolutionary trajectories within PWH.

A central finding of our work is that variation in HIV-1 LTR activity across isolates and clades is substantial and cannot be inferred reliably from phylogenetic grouping alone. Previous studies reported differences in LTR activity between select clades—most notably the greater activity of clade C attributed to additional NF-κB sites—but were limited to a handful of isolates.^36,37^ Our large-scale functional analysis confirms these trends while revealing extensive intra-clade and within-PWH heterogeneity driven by differences in TF configurations. These TF configurations not only influence baseline and signal-responsive transcriptional activity but also determine robustness across primary CD4+ T cell donors, suggesting that certain regulatory configurations buffer host-to-host variation in signaling environments. This is consistent with recent observations that proviral sequence features contribute to heterogeneous inducibility across reservoirs.^3,50,80,81,82^ Together, our results have important implications for interpreting viral fitness, latency dynamics, and responsiveness to immune signaling across genetically diverse reservoirs and hosts. In particular, differences in regulatory architectures, which result in differential TF and cofactor recruitment, suggest that latency reversal agents may not uniformly reactivate non-defective proviruses even when considering integration site effects, providing a potential explanation for the incomplete efficacy of current shock-and-kill strategies.^17,50,83^

Our findings further highlight a previously underappreciated regulatory compensation within retroviral genomes. The negative correlation between HIV-1 U3 and U5 activity across isolates suggests that distinct LTR regions can offset one another to maintain balanced transcriptional output. This compensation may also be associated with different clinical outcomes. For instance, 01_AE LTRs have a weak response to TNFα and CD3 stimulation in U3 but a strong response to IFNγ in U5. Clade 01_AE more frequently exhibits CXCR4 tropism than clades B or C, enabling infection of naïve and central memory CD4+ T cells which are essential for maintaining the CD4+ pool and are characterized by low baseline NF-κB activity.^84,85,86^ In IFN-rich lymphoid environments, reduced reliance on NF-κB-driven activation coupled with enhanced sensitivity to STAT/IRF signaling may sustain viral transcription within these renewal compartments. This, in turn, could promote preferential depletion of naïve and central memory CD4+ T cells, contributing to the more rapid immunologic decline and accelerated disease progression observed in 01_AE-infected cohorts.^87,88^

Transcriptional compensation extends beyond the LTR. Differences in regulatory logic between HIV-1 and HIV-2 LTRs, such as the absence of IRF-dependent regulation in the HIV-2 LTR, are partially compensated by intragenic CREs proximal to the LTR. Likewise, the relative scarcity of ETS sites in most HIV-1 LTRs, present in the HIV-2 LTR, is complemented by ETS sites within the HIV-1 CRE in *gag*. Together, this supports a model in which HIV transcription is governed by modular and distributed regulatory architectures that complement the main promoter region.

The systematic identification of intragenic CREs adds an important layer to HIV regulatory variation. While a subset of intragenic CREs in HIV-1 have been described previously,^18,19,20,21,22^ our work demonstrates that such elements are widespread across HIV-1 isolates and are also present in HIV-2, with some being evolutionarily conserved and others more variable. A subset of intragenic CREs have potential functional roles in HIV transcription, including the CRE in HIV-1 *gag* which is under evolutionary constraint. Others, however, may be a byproduct of selective pressures acting on viral protein-coding sequences. For instance, we observe extensive variation in intragenic CREs across HIV-2 isolates, arguing against a conserved functional role for these elements. In addition, our longitudinal analyses show a lack of directional selection for transcriptional output in the CRE in HIV-1 *env,* but rather activity variation likely reflecting neutral evolution or selection acting on protein-coding constraints. Regardless of whether they impact viral replication and gene transcription, these intragenic CREs may result in the production of pathogenic RNAs or proteins. Intragenic transcription in HIV-1 has been observed in individuals on suppressive ART and linked to immune activation and dysfunction.^22,67,68^ Our discovery of multiple conserved CREs capable of driving such transcription suggests that persistent viral RNA production may be frequent in proviral genomes rather than a rare aberration. Because most proviruses in ART-treated individuals are defective, this raises the possibility that even LTR-defective proviruses may remain transcriptionally active through alternative regulatory elements, contributing to the chronic inflammation observed in PWH.^50,51,89^ Defining the functional consequences of intragenic CREs and determining how their regulatory architectures relate to clinical phenotypes will be important directions for future studies.

A key advance of our study is the development of models that predict transcriptional activity from sequence. CREST accurately predicts baseline transcriptional activity in Jurkat cells for any DNA sequence, which enables activity predictions across genomic regions and isolates allowing the identification of novel and low-conserved CREs, and recapitulates motif-level contributions. LARM, instead, predicts stimulus-induced activation and captures reactivation in integrated proviral contexts despite being trained on episomal MPRA data. Together, these models provide a scalable framework for interpreting new viral sequences, longitudinal evolution, and datasets derived from PWH without requiring exhaustive experimental assays. This predictive capacity opens the door to functional annotation of viral diversity at a scale comparable to genomic surveillance efforts. We provide open-source code for large-scale batch analysis of user-provided sequences, along with a web-based resource for exploring our datasets, model predictions, and user sequence submissions.

Together, our results provide a framework for linking viral sequence variation to transcriptional behavior across isolates and within individuals, with implications for reservoir heterogeneity and responses to reactivation strategies. More broadly, this work demonstrates that principles of cis-regulatory grammar, modularity, and evolutionary constraint operate even within compact viral genomes, highlighting how complex regulatory programs can emerge from minimal genetic systems.

### Limitations of this study

While our approach enables systematic dissection of HIV regulatory architecture at scale, several considerations frame the interpretation and extension of these findings. MPRA measurements were performed largely in episomal contexts, which resolve intrinsic regulatory potential and motif-level contributions but do not capture integrated proviral environments or site-specific effects known to influence HIV transcription and latency. Notably, both MPRA-derived activation responses and LARM predictions correlated with reactivation of integrated proviruses, indicating that our framework captures meaningful population-level regulatory behaviors despite lacking locus-specific resolution.

The use of ∼200 bp tiles enabled fine-scale mapping of regulatory modules but may miss longer-range interactions among TF binding sites across extended LTR regions. Future approaches incorporating longer regulatory fragments or integrated systems may further refine higher-order regulatory logic, and consider the influence of HIV Tat on LTR regulation.^90^

Our analyses focused primarily on Jurkat cells and primary CD4+ T cells, whereas HIV reservoirs span diverse cellular compartments, including distinct T cell subsets, myeloid cells, and tissue environments with unique TF landscapes. Extending MPRA profiling and predictive modeling into these contexts will provide additional insight into how cis-regulatory architectures interact with host-specific signaling environments.

## Resource availability

### Lead contact

Further information and requests for resources and reagents should be directed to and will be fulfilled by the lead contact, Juan Fuxman Bass.

## Materials availability

This study did not generate new unique reagents. All materials generated in this study are available from the lead contact with a completed materials transfer agreement.

## Data and code availability

All data are included in the main figures and supplemental figures/tables.

This paper reports original code. All custom analysis code is publicly available at https://github.com/FuxmanBass-lab/HIV-MPRA, https://github.com/FuxmanBass-lab/MPRA, and https://github.com/osyafinkelberg/mpra-predictor, and is listed in the Key Resources Table. Any additional information required to reanalyze the data reported in this paper is available from the lead contact upon request.

## Acknowledgements

We thank Dr. Todd Blute for assistance with flow cytometry as well as Mr. Brian Tilton, BUMC Flow Cytometry Core Facility, and Yun Shen for IT assistance installing and adapting packages in the computer cluster. Also, assistance was provided by Providence/Boston CFAR Scientific Core (P30 AI042853). This work was supported by National Institutes of Health grants R35GM128625 (to J.I.F.B.), R35HG011329 (to R.T.), R01 AI187175 and DA055488 (to A.J.H.), R01AI151051 (to T.S.), DP2AI183504 (J.P.R.); U01AI176320 (to J.P.R.); R01DK140972 (to J.P.R); and Crohn’s & Colitis Foundation grant 1158945 (to J.P.R.). J.A.F. and M.Y.E were supported by NIH grant T32GM150533. M.Y.E. was supported by NIH grant T32GM154655 and D.L.B. was supported by T32AI052074. E.M. was supported by the National Science Foundation grant BIO-1659605. J.P. was supported by Boston University’s undergraduate research opportunities program.

## Author contributions

Conceptualization, J.I.F.B., B.E., M.Y.E., J.A.F., T.T.; methodology, B.E., M.Y.E., S.K., J.A.F., T.T., D.L.B., L.S-U., J.I.F.B., R.T., T.S., A.J.H., J.P.R.; software, J.A.F., M.Y.E., L.S-U., G.M-E., R.T., R.C.; validation, B.E., T.T., M.Y.E., J.A.F., R.T., D.L.B.; formal analysis, B.E., M.Y.E., J.A.F., L.S-U., R.C., H.C., M.D., J.I.F.B., T.S., R.T.; investigation, B.E., T.T., D.L.B., S.K., M.D., E.M., J.P., B.D., C-H.H.; resources, J.I.F.B., R.T., T.S., A.J.H., J.P.R.; data curation, B.E., T.T., J.A.F., M.A.P., D.L.B., J.I.F.B.; writing – original draft, J.I.F.B., B.E., M.Y.E., J.A.F., T.T.; writing – review and editing, J.I.F.B., B.E., M.Y.E., J.A.F., T.T., A.J.H., R.T., T.S., J.P.R., L.S-U., D.L.B.; visualization, J.I.F.B., B.E., M.Y.E., J.A.F., T.T., D.L.B., M.D.; supervision, J.I.F.B., R.T., T.S., J.P.R., A.J.H.; project administration, J.I.F.B., R.T; and funding acquisition, J.I.F.B., R.T., T.S., J.P.R., A.J.H.

## Declaration of interests

The authors declare no competing interests.

## Declaration of generative AI and AI-assisted technologies in the writing process

During the preparation of this work, the author(s) used ChatGPT to improve clarity, grammar, and academic writing style. After using this tool or service, the author(s) reviewed and edited the content as needed and take full responsibility for the content of the publication.

## STAR Methods

### MPRA viral tiling library sequence design

Viral sequences used for generating the MPRA tiling library are listed in **Table S1**, including virus and strain, accession number, and sequence length. Each virus was tiled using 200 bp sequences with 50 bp offsets and each tile was tested in both orientations relative to the minimal promoter in the MPRA vector. Tile generation begins from the 5’ end of the sequence provided for each virus and the offset applies until the 3’ end of the sequence. To avoid having tiles shorter than 200 bp, the last tile for each virus corresponds to the last 200 bp sequence from the 3’ end. Overall, we generated 68,612 tile sequences for the viruses listed.

In addition, we included negative and positive control sequences in the MPRA library to normalize activity for experimental tiles and to evaluate the sensitivity of each MPRA experiment. Negative controls consisted of 250 shuffle sequences, 1,876 sequences overlapping human ORFs, and 506 sequences without detected MPRA activity in prior studies. The positive control set of tiles consisted of 91 human genome sequences known to be active in MPRA, representing a broad activity spectrum. These control sequences correspond to those used in our previous studies.^26,27^

### MPRA saturation mutagenesis and isolate library design

The saturation mutagenesis and isolate library was generated as previously described.^27^ Briefly, for saturation mutagenesis, we selected 10 tiles with strong CRE activity (**Table S2**) from HIV-1 ReJo (Genbank: JN944911.1) and HIV-2 ROD (GenBank: M15390.1). To minimize redundancy, we avoided selecting tiles that overlapped by more than 100 bp. Saturation mutagenesis was performed from positions 6 to 195 of each selected 200 bp tile by introducing all possible single-nucleotide substitutions, wherein each nucleotide position was individually mutated to the other three possible bases, resulting in a total of 570 unique single-nucleotide variants per tile tested.

For the tiles indicated above, we have also evaluated homologous sequences corresponding to isolates from the same viral species. In June 2023, we retrieved all publicly available complete genome assemblies for HIV-1 and HIV-2 from the NCBI Virus portal, applying the “complete” filter to exclude partial or draft records and recording the download date to ensure reproducibility (**Table S3-4**). Using Python and Biopython’s local alignment mode, we sliced each sequence into 200-nt tiles (including their reverse complements) and aligned them against every reference genome with a scoring scheme of +2 for matches, 0 for mismatches, and gap penalties of −20 (opening) and −2 (extension). From each alignment we extracted the aligned length and the number of exact matches for both forward and reverse orientations, then merged these results into a unified dataset for downstream analysis.

Next, we imported the alignment outputs into R and performed a global realignment using Biostrings’ pairwiseAlignment with a nucleotide substitution matrix (match = 2, mismatch = −3), a gap opening penalty of −5, and a gap extension penalty of −2. We filtered out any alignments shorter than 180 nt or containing fewer than 100 exact matches. To guard against overly extended outliers, we calculated the 95th percentile of the remaining alignment lengths for each tile and set a dynamic maximum length threshold: tiles whose lengths exceeded this threshold (removing at most the top 5 % of extensions) were discarded, while the remainder were retained. Tiles identified as originating from reverse-strand alignments were reverse-complemented so that all sequences share a uniform 5′→3′ orientation. In a final pass, any sequence still longer than 200 nt was trimmed to its last 200 nt, its start position adjusted accordingly, and its exact-match count recalculated via one more global alignment. This procedure yields a final collection of 180-200 nt tiles that uniformly satisfy our stringent length and match criteria. This library also included the same set of positive and negative control sequences as the tiling library above.

### MPRA library construction

The two MPRA libraries were constructed as previously described.^25,27^ Briefly, oligonucleotides were synthesized (Twist Biosciences) in the range of 210 to 230bp sequences containing 180 to 200bp of viral genomic sequences and 15bp adaptor sequence on both ends. Unique 20bp barcodes were added by PCR into a backbone vector (addgene #109035) by Gibson Assembly. The Gibson assembly product was electroporated into NEB 10-beta E.coli, and the resulting plasmid library was sequenced by Illumina 2x150bp chemistry to determine oligonucleotide-barcode pairings. The library then underwent restriction digest using AsiSI, and GFP with a minimal TATA promoter was inserted by Gibson assembly resulting in the 180-200bp oligo sequence positioned directly upstream of the promoter and the 20bp barcode located in the 3’ UTR of the GFP. After library expansion in *E. coli*, the final MPRA plasmid library was sequenced by Illumina 1x26bp chemistry to acquire a baseline representation of each oligo-barcode pair within the library.

### MPRA library transfection into Jurkat cells

Jurkat cells (ATCC-TIB-15) were cultured in RPMI media (Fisher Scientific, Catalogue # A1049101) with 10% Fetal Bovine Serum (R&D Systems, Catalog # S12450H) and 1% Antibiotic-antimicotic (Thermofisher Scientific, Catalogue #15240062) up to a density of 1 million cells per mL prior to transfection. For each replicate, 100 million cells were electroporated with 100 µg of MPRA library plasmid using the 100 µL Neon Transfection System kit with 3 pulses of 1350V for 10 ms. Cells were then incubated at 37 °C for 24 hours in the corresponding media without antibiotic-antimicotic to allow for expression of the barcoded reporter construct. Cells were then left unstimulated or were stimulated with 20 ng/mL of IFNγ (Invivogen cat# rcyec-hifng) for 6 hours, 20 ng/mL of TNFα (Invivogen cat# rcyc-htnfa) for 1 hour, or 10 ng/mL Phorbol 12-myristate 13-acetate (Millipore Sigma, Cat# P8139-1MG) and 1ug/mL Anti-CD3 (Fisher Scientific, Cat# 16-0037-8**5**) for 1 hour. Finally, cells were pelleted, washed three times with PBS, and stored at –80 °C for later RNA extraction. Experiments were performed in biological quintuplicates.

### MPRA library transfection into human primary CD4+ T cells

Peripheral blood mononuclear cells (PBMCs) were isolated from fresh apheresis leukoreduction packs (BloodWorks) obtained from four fully deidentified donors using Ficoll-Paque Plus (GE, 17-1440-03). CD4+ T cells were subsequently enriched via magnetic isolation (BioLegend, 480130) and maintained in CTS OpTmizer T cell Expansion Medium (ThermoFisher, A1048501) supplemented with 5% FBS, 1% glutamine, 100 U/mL penicillin-streptomycin, and 55 mM 2-mercaptoethanol. Approximately 100 million CD4+ T cells were activated using human T cell activation beads (Miltenyi, 130-091-441) and recombinant human IL-2 (100 U/mL; NCI Biological Resources Branch) for 48–72 hours.

Cells were transfected using Neon NxT electroporation system (ThermoFisher # N10096). In brief, 100 million activated cells from four separate donors were counted, washed twice in PBS, and resuspended in 1 mL of R buffer and then each mixed separately with 100 μg of MPRA library plasmid. The mixture from each donor was divided into 10 reactions and electroporated with the program (1600 volt, 10 ms, 3 pulses). After electroporation, cells were transferred to 50 mL of prewarmed media in T75 flasks immediately and then incubated at 37°C for 24 hours. Right before harvesting the cells, transfection efficiency was checked according to flow cytometry, finding 34.7-65.8% GFP+ cells for all donors. Cells were then collected by centrifugation, washed once with PBS, collected and frozen at −80°C.

### RNA extraction and MPRA RNA-seq library preparation

The MPRA RNA library was prepared as previously described.^25^ Briefly, RNA was extracted from frozen cell pellets using the Qiagen RNeasy Maxi kit (Cat# 75162). Half of the isolated total RNA underwent DNase treatment and a mixture of three GFP-specific biotinylated primers (#120, #123 and #126)(**Table S11**) were used to capture GFP transcripts with Streptavidin C1 Dynabeads (Life Technologies, cat# 65002). An additional DNase treatment was performed to eliminate residual RNA. cDNA was synthesized from GFP mRNA with primer #19 using SuperScript III (Thermo, Cat# 18-080-400) and purified with AMPure XP beads (Beckman Coulter Life Sciences, Cat#A63880).

To minimize amplification bias during the creation of cDNA tag sequencing libraries, samples were amplified by qPCR to estimate relative concentrations of GFP cDNA using 1 μL of sample in a 10 μL PCR reaction containing 5 μL Q5 NEBNext master mix, 1.7 μL SYBR Green I diluted 1:10,000 (Life Technologies, S-7567) using primers specific for the GFP transcript (#781 and #782)(**Table S11**). Samples were amplified with the following qPCR conditions: 95 °C for 20 s, 40 cycles (95 °C for 20 s, 65 °C for 20 s, 72 °C for 30 s), 72 °C for 2 min. The optimal cycle number for final library amplification was defined as n-1, where n represents the threshold cycle (Ct) determined by qPCR. Replicates within each condition were diluted to the same concentration based on the qPCR results.

Illumina sequencing libraries were constructed using a two-step amplification process to add sequencing adapters and indices. An initial PCR amplification with NEBNext Ultra II Q5 Master Mix (NEB, Cat#M0544X) and primers #781 and #782 were used to extend adapters. To minimize over-amplification during library construction, the number of PCR cycles used in the first amplification was selected based on the qPCR above and required 8 to 11 cycles in this experiment. A second 6 cycle PCR using NEBNext Ultra II Q5 Master Mix added P7 and P5 indices and flow cell adapters (**Table S11**). Briefly, 10 μL of cDNA samples and mpra:minP:gfp plasmid control (diluted to the qPCR cycle range of the samples) were amplified using the reaction conditions from the qPCR scaled to 50 μL, excluding SYBR Green I. Amplified cDNA was SPRI purified and eluted in 40 μL of EB. Individual sequencing barcodes were added to each sample by amplifying the entire 40 μL elution in a 100 μL Q5 NEBNext reaction with 0.5 μM of TruSeq_Universal_Adapter primer and a reverse primer containing a unique 8 bp index (Illumina_Multiplex, CAAGCAGAAGACGGCATACGAGATXXXXXXXXGTGACTGGAGTTCAGACGTGTGC) for sample demultiplexing post-sequencing. Samples were amplified at 95 °C for 20 s, six cycles (95 °C for 20 s, 64 °C for 30 s, 72 °C for 30 s), 72 °C for 2 min. Indexed libraries were purified with 1.8X SPRI and pooled according to molar estimates from BioAnalyzer quantifications. The resulting MPRA RNA-tag libraries were sequenced using paired-end Illumina chemistry and dual index reads on either a Illumina Sequencing PE150, NextSeq 1000 or NovaSeq 6000 instrument.

### MPRA data analysis

MPRA sequencing data were processed using a custom pipeline adapted from MPRASuite^25,91^ (https://github.com/tewhey-lab/MPRASuite) and rewritten in Python/R/shell (https://github.com/FuxmanBass-lab/MPRA). The workflow comprised four stages: barcode–oligonucleotide matching, barcode counting, activity modeling, and condition comparisons.

Barcode–oligonucleotide matching: Paired-end reads from the cloned plasmid library were used to build a barcode–oligonucleotide dictionary by extracting barcodes and aligning oligonucleotide sequences to the reference library FASTA. While mismatches were allowed when aligning the MPRA tiling library (0.05 mismatch rate), alignments for the MPRA isolates and saturation mutagenesis library required exact matches for the MPRA isolates and saturation mutagenesis library, to account for single-base differences in the library design. Only high-confidence, uniquely mapped barcode–oligonucleotide pairs were retained. Library QC included barcode complexity per oligonucleotide and alignment/error metrics.

Counting and aggregation: For each replicate, RNA (and plasmid DNA) reads were parsed to count barcodes, which were then assigned to oligonucleotides using the barcode–oligonucleotide dictionary to generate barcode and oligonucleotide-level count matrices. Oligonucleotides with insufficient barcode support were excluded. Replicate-level QC included recovered barcodes per oligonucleotide and sequencing depth.

Activity modeling and normalization: Oligonucleotide activity was inferred in DESeq2^92^ by fitting a negative binomial generalized linear model to joint DNA and RNA count matrices with DNA as the reference. Size factors were first estimated by DESeq2’s median-of-ratios method, then DNA libraries were anchored by forcing DNA size factors to 1 and rescaling RNA size factors by the median DNA size factor, so RNA signals reflect expression per plasmid input. For each RNA condition, we then applied summit-shift normalization by adjusting the condition-specific size factor so that the mode of the log₂(RNA/DNA) fold-change distribution computed on negative control oligonucleotides was centered at 0. Finally, dispersions were estimated with a local fit and then replaced with cell-type-specific dispersion estimates to capture condition-dependent variance.

Comparison modeling: For each condition comparison, differential activity was tested with DESeq2 on RNA counts using a negative binomial generalized linear model with a plasmid baseline offset (log mean plasmid counts per oligonucleotide). Dispersions were estimated with a local fit.

DESeq2 MPRA data were further analyzed across experimental conditions after filtering for tiles with sufficient DNA representation (≥50 DNA counts per tile)(**Tables S1**, **S2**, and **S4**). The ≥50 DNA counts threshold was chosen based on activity variance analysis, where activity estimates became largely independent of DNA counts above this level.

### MPRA quality control and data reproducibility

To assess MPRA library quality and reproducibility, we evaluated barcode representation per oligonucleotide and concordance across experimental conditions. In the tiling library, the distribution of unique barcodes per oligonucleotide was consistent across plasmid, Jurkat, and K562 samples (**Figure S1A**). Similarly, in the saturation mutagenesis and isolates library, barcode representation was comparable across plasmid and all stimulation conditions (unstimulated, αCD3 + PMA, TNFα, and IFNγ; **Figure S1B**), as well as in primary CD4⁺ T cells (**Figure S1F**).

We observed high reproducibility of normalized oligonucleotide counts across biological replicates and conditions in both libraries (**Figures S1C-D** and **S1G**). In addition, positive control tiles showed highly concordant activity between unstimulated and stimulated conditions (αCD3 + PMA, TNFα, and IFNγ; **Fig. S1E**), indicating that global stimulation does not systematically alter control element activity. Together, these metrics support robust barcode representation, high reproducibility, and consistent activity measurements across conditions.

### Saturation mutagenesis analysis

Saturation mutagenesis variants were analyzed to quantify the MPRA activity impact of each single–nucleotide substitution within a subset of transcriptionally active genomic tiles (**Table S2**). The analysis pipeline is available in github (https://github.com/FuxmanBass-lab/HIV-MPRA/tree/main/Satmut). Briefly, for each tile, all single–nucleotide variants were grouped with their matched reference sequence. Variant activity was quantified using MPRA-derived log₂(RNA/DNA) fold changes, and per-mutation effects were computed as the difference in activity between each variant and its matched reference sequence.

To stabilize estimates across dense variant sets within each tile, mutation effect sizes were modeled using an empirical Bayes framework that shrinks variant activities toward a tile- and cell-type–specific baseline. This approach accounts for heteroskedastic uncertainty across variants while preserving relative effect sizes. For each variant, posterior-shrunken effect estimates were used to compute mutation-specific skew statistics, and statistical significance was assessed using z-tests with false discovery rate correction performed independently for each tile, condition, and cell type.

Mutation effects were visualized using hybrid sequence-logo and lollipop plots. At each nucleotide position, the reference base was represented as a sequence-logo letter with height proportional to its estimated effect, while alternate alleles were shown as lollipops indicating the magnitude and direction of their effects. Plots were generated independently for each cell type and stimulation condition using symmetric scaling to facilitate direct comparison.

### Generation of full-length single-round HIV-1 clones with variant LTRs

Variant LTR-HIV-ΔEnv-EGFP constructs were generated using PCR, overlap extension PCR, restriction digests, and Gibson and HI-Fi assembly (NEB) to replace the 3’-LTR in NL43-ΔEnv-EGFP (Aids reagent program, ARP-11100) with LTRs from the indicated virus were derived from sequences in the Los Alamos database. In some instances, full length proviral LTRs were generated by aligning the genomic LTRs and combining the U3, R, and U5 regions through part of Psi. The LTR sequences used are shown in **Table S12** along with their corresponding accession number from the Los Alamos database. The resulting plasmids were validated using nanopore sequencing from Plasmidsaurus. The LTRs were cloned into the 3’ LTR to allow for near equivalent virus production from HEK293T cells.^93^ Once infection in a target cell occurs, reverse transcription will lead to the U3 region of the 3’ LTR, containing the core promoter/enhancer region becoming the U3 region of the 5’ LTR.

### Virus production and infection

VSVg-pseudotyped viruses were created through PEI co-transfection in HEK293T cells with a plasmid expressing the VSV glycoprotein as previously described.^94^ Viral containing supernatants were collected and concentrated by centrifugation through a 20% sucrose gradient at 56,000 xg for 1.5 hours in a vacuum. Viral pellets were resuspended for 24 hours in media then aliquoted and stored at -80 °C for future use.

To determine multiplicity of infection (MOI), unmodified NL43-ΔEnv-EGFP was titrated in Jurkat T-cells using flow cytometry and amounts of p24 were determined using an ELISA. Equivalent amounts of p24 from each viral prep were added to Jurkat T-cells equating to an MOI of 0.5 for NL43-ΔEnv-EGFP. The cells were spinoculated at 1200 xg for 1.5 hours at room temperature, then incubated at 37 °C in 5% CO2 for at least 1 hour. The virus containing media was removed, and cells were washed 1x in media then resuspended at a final cellular concentration of 1 million cells per mL of medium and cultured at 37 C, 5% CO2 in RPMI supplemented with 10 % (v/v) FBS, L-glutamine, and penicillin/streptomycin (R10) for 48 hours.

### Fluorescence activated cell sorting (FACS) of infected cells and reactivation

Infected cells were collected via centrifugation and resuspended in PBS with 2% FBS and 1 mM EDTA (FACS buffer) containing a 10x concentration of Calcein Blue AM (ThermoFisher, C34853) for at least 15 min prior to sorting. Cells were sorted on a MoFlow Astros with the assistance of the Boston University Flow Cytometry Core Facility (BU FCCF) for GFP+ and GFP- signals and collected in RPMI supplemented with 20 % (v/v) FBS, L-glutamine, and penicillin/streptomycin. Cells were washed, then cultured in R10 for seven days with increasing medium every 2-3 days. On day 7, the cells were reactivated with 20 ng/mL TNFα (PeproTech, 300-01A), 10 ng/mL PMA/ 1 μg/mL anti-CD3 (Biolegend, 317302, Millipore Sigma, P1585, Invivogen,tlrl-pma), or 0.1 % (v/v) DMSO (Vehicle) for 48 h. Reactivated cells were harvested, stained with Live/Dead fixable blue in PBS for 30 min, fixed with 4 % methanol-free PFA for 15 min, and resuspended in FACS buffer with a PBS wash between each step. Flow cytometry was done using a Cytek Aurora through the BU FCCF. Each sample was gated on the uninfected control plus the indicated stimuli for each replicate (**Figures S3F-G**). The fold-change in reactivation relative to DMSO control and geometric mean fluorescence intensity (MFI) were determined, and the correlation of these parameters to the measured MPRA activity for tile HXB2:250-431 or the activity predicted by LARM were calculated.

### Motif grammar analysis in HIV-1 sequences

To characterize TF motif grammar within HIV-1 tiles, we performed systematic motif scanning and combinatorial analysis across viral isolates (https://github.com/FuxmanBass-lab/HIV-MPRA/tree/main/Motifs). Position weight matrices (PWMs) for human TFs were obtained from the JASPAR 2024 CORE database,^95^ requiring a minimum information content of 10, and were further filtered to a curated TF panel based on TF motif family membership. Motif scoring was performed using log-odds position-specific scoring matrices with a fixed viral genome–wide nucleotide background estimated across all analyzed isolates. Motif instances were scanned on both strands using FIMO, and candidate hits were required to satisfy uniform score and significance thresholds (FIMO score ≥10, p ≤ 1×10⁻⁴, q ≤ 0.10)(**Table S6**). Only high-confidence motif matches passing these criteria were retained for downstream analysis.

Motif instances were then deduplicated and assigned in a hierarchical, precedence-aware manner to yield biologically interpretable site calls. Overlapping or near-duplicate hits arising from multiple PWMs corresponding to the same TF were first collapsed using non-maximum suppression, prioritizing higher-scoring motif instances. TF-level hits were mapped to TF motif families, with co-occurring motifs from multiple constituent families collapsed into composite meta-families where applicable. Finally, overlapping motif instances across different TF motif families were resolved using a locus-level exclusivity scheme that enforced a single TF motif family assignment per genomic position based on predefined family precedence, motif strength, and positional criteria.

To compare TF motif usage across HIV-1 clades, motif counts per tile were aggregated at the isolate level (**Table S6**). HIV-1 isolates were grouped into major clades using curated subtype annotations. For a set of TF motif families, motif counts were summarized within each clade by computing the distribution of motif counts across isolates. For each tile, these distributions were visualized as a matrix of pie charts, with rows corresponding to clades and columns to TF motif families, enabling direct comparison of motif abundance patterns across clades.

To resolve differences in motif organization within HXB2:250-431, motif hits for the NFKB/REL and SP/KLF families were first analyzed in a common coordinate system. Starting from the per-isolate motif hit table, we retained deduplicated sites and mapped each hit from isolate-specific coordinates onto a shared reference axis using per-isolate global alignments (**Figure S4A**). This coordinate “lift-over” enables direct comparison of motif positions across divergent HIV-1 sequences. We then summarized positional preference for each TF family by plotting normalized hit-density profiles across the tile, complemented by a range-based coverage track that captures how often each base position is spanned by motif instances. Building on these aligned coordinates, we discretized the tile into canonical positional bins capturing recurring NFKB/REL and SP/KLF site windows. For each isolate, we recorded the presence/absence of motif hits in each of the four NFKB/REL and four SP/KLF bins and encoded this as a compact motif signature. These isolate-level grammar groups were then linked to MPRA activity measurements: for each grammar with sufficient isolate support (≥3 isolates), we visualized the distribution of CD4+ T cells or Jurkat baseline MPRA activity (log2 fold-change) or activation by αCD3 + PMA, or TNFα across isolates as violin plots, alongside a schematic glyph of the underlying site pattern and a clade-distribution pie chart. This integrates sequence architecture, lineage composition, and functional output in a single view, allowing specific positional motif configurations to be compared for both prevalence and regulatory effect.

For tile HXB2:581-780, the same reference-aligned analysis framework was applied using a tile-specific TF set. Motif hits for IRF, E2F, and SP/KLF were mapped to a shared reference coordinate system and discretized into predefined positional bins (**Figure S4C**). These bins were used to encode isolate-level motif grammars, which were associated with clade composition, CD4+ T cell and Jurkat baseline MPRA activity, and IFNγ activation.

In primary CD4+ T cells, donor-to-donor variability in MPRA activity was quantified across four donors. For each tile, log2 MPRA activity was summarized across donors, and variability was modeled as a function of mean activity by fitting a global mean–variance trend on a deduplicated tile set. Deviations from this trend were quantified using an F-like statistic, defined as the ratio of observed variance to the variance expected from the fitted trend, and linked to isolate groups defined by motif grammar signatures and clade annotations.

### Variation in MPRA baseline activity and activation

To prevent low-activity tiles (0–1 in linear scale) from disproportionately influencing the results, a pseudocount of +1 in linear scale was added symmetrically to both control and treatment replicate activities derived from DESeq2 normalization prior to log₂ transformation. The resulting pseudocount added log₂FC values were used for plotting violins and dot heatmaps. Subclades A1, A2, A4, and A6 were consolidated under major clade A, and subclades F1, F2, and F1F2 were consolidated under major clade F. The distribution of MPRA activity (log₂(fold activation)) in Jurkat and CD4+ T cells is shown as violin plots across isolates from different viruses and clades as indicated in the corresponding figures. Dot heatmaps were generated for each tile, with rows corresponding to activation conditions (αCD3+PMA, TNFα, or IFNγ) and columns corresponding to clades. Dot color encodes the median log₂(fold activation) using a diverging scale centered at zero, while dot size encodes variability defined as the difference between the 95th and 5th percentiles (P95–P5) of log₂(fold activation) values. To enable comparisons across tiles and conditions, dot radii were scaled linearly using a fixed global denominator (P95–P5 = 2.0) rather than tile-dependent normalization.

### Variation in MPRA activity across recombinant clade isolates

To determine the parental clade origin at region HXB2:250-431 in recombinant strains, the isolate sequence at this region from each recombinant isolate was aligned to the HIV-1 reference genome HXB2 (**Table S5**). The resulting HXB2 coordinates were intersected with Circulating Recombinant Form (CRF) breakpoint annotations from the Los Alamos HIV Sequence Database (https://www.hiv.lanl.gov), enabling assignment of which parental clade contributed the LTR segment corresponding to tile HXB2:250-431. For recombinant strains in which the tile HXB2:250-431 is assigned to a parental clade based on breakpoint overlap, the corresponding accessions were retained for downstream analysis. Dot heatmaps were generated as above, using only recombinant clades represented by at least three independent isolates.

### MPRA activity variation from isolates within PWH

HIV-1 isolates corresponding to MPRA-tested tiles were retrieved from the Los Alamos HIV Sequence Database (https://www.hiv.lanl.gov). PWH identifiers and codes were obtained from Los Alamos metadata whenever available. For accessions lacking PWH-level annotation in Los Alamos, information was curated from the original source publications referenced in the associated GenBank records (**Table S3**).

Analyses were performed for HIV-1 tiles HXB2:250-431, HXB2:581-780, HXB2:1330-1530 and HXB2:7784-7984 (**Table S4**). Only isolates with activity measurements across all shown baseline or stimulation conditions were retained. Proviral DNA–derived sequences and viral RNA-derived sequences were processed and analyzed separately. For each tile, PWH were included only if at least five isolates were available. For each PWH and tile, the distribution of activity or fold activation is plotted as well as the distribution of TF configurations.

### Sequence alignments of isolates within PWH

Within each PWH and tile, identical tile sequences were collapsed to unique sequence variants while retaining the number of occurrences of each variant to capture within-PWH sequence frequency. Unique sequence variants were aligned using MAFFT^96^ with automatic parameter selection to generate multiple sequence alignments that preserve positional homology across isolates. Alignments were performed independently for each PWH and tile to ensure that sequence variation was evaluated strictly within a PWH-specific evolutionary context. Aligned sequences served as the reference framework for downstream visualization and analysis, allowing polymorphic sites, conserved regions, and indel structure to be directly compared alongside MPRA-derived regulatory activity measurements across unstimulated and stimulated conditions.

### Codon- and amino-acid–level diversity profiling across HIV-1 Gag

We analyzed a Los Alamos National Laboratory HIV-1 pre-aligned nucleotide multiple sequence alignment (MSA) of Gag isolates that includes the HXB2 reference sequence. HXB2 genomic coordinates were assigned by traversing the ungapped HXB2 sequence within the alignment, such that each non-gap alignment column in HXB2 was mapped to its corresponding HXB2 coordinate, whereas alignment columns corresponding to gaps in HXB2 were excluded from coordinate assignment. Codon positions were then defined in the HXB2 Gag reading frame by stepping through the coding sequence in groups of three nucleotides beginning at the Gag start codon.

For codon-level analyses, we considered only codons for which all three nucleotide positions were present in the HXB2 coordinate map, thereby ensuring complete codon representation in the aligned region. Entropy calculations were performed across non-reference isolates only, excluding HXB2 from the distributions. For each codon, isolates contributed only if all three aligned nucleotide positions contained unambiguous bases (A, C, G, or T); codons containing gaps or ambiguous bases in a given isolate were excluded for that isolate. Codons with fewer than 50 usable non-reference isolates were omitted from downstream entropy analyses. For each eligible codon, we computed Shannon entropy over observed codons, Shannon entropy over the encoded amino-acid distribution, and conditional codon entropy given amino acid, H(codon | a.a.), to distinguish total codon diversity from amino-acid–changing and amino-acid–preserving components of variation (https://github.com/FuxmanBass-lab/HIV-MPRA/tree/main/Entropy).

To test whether the focal Gag region showed reduced diversity relative to the surrounding coding sequence, we performed a motif-independent window-based depletion analysis on these codon-level entropy tracks. A region of interest was defined in HXB2 coordinates, and codons whose HXB2-mapped intervals overlapped this window were classified as in-window. For each metric, we calculated Δ median as the median value within the window minus the median value outside the window. Statistical significance was assessed using a one-sided circular-shift permutation test over codon positions (10,000 permutations), in which the same-sized window mask was rotated along the codon series to generate a null distribution while preserving window length and contiguity. P values therefore test whether the observed window exhibits unusually low entropy relative to the remainder of the analyzed Gag region.

### CREST model architecture

We developed CREST (Cis-Regulatory Element Sequence Tester), a deep learning model that predicts MPRA activity in Jurkat cells from 200 bp DNA sequences (https://github.com/osyafinkelberg/mpra-predictor). The available experimental dataset, generated previously,^27^ consists of 65,028 unique tile sequences corresponding to viral sequences, and positive and negative control sequences derived from the human genome. This dataset size is approximately an order of magnitude smaller than those typically used to train deep neural network MPRA predictors.^26,97^ Therefore, we employed a transfer-learning approach.

We selected two pretrained source models, Malinois^26^ and Sei^54^, which capture sequence features relevant to Jurkat MPRA activity while accepting input sequence lengths comparable to the MPRA plasmid. This choice was motivated by three considerations: (i) MPRA activity is strongly correlated between Jurkat and K562 cells; (ii) Malinois, which we previously developed, predicts MPRA activity in K562 with good accuracy; and (iii) sequence binding preferences of Jurkat-specific transcriptional regulators are expected to be well captured by the Sei model.

We fully transferred the convolutional component of the Malinois architecture, consisting of three convolutional layers with batch normalization and max pooling, as well as the convolutional layers of the Sei model preceding the spline pooler. The 200 bp input tile sequence was flanked with constant MPRA plasmid sequences to match the input format expected by both models. All weights of the Malinois and Sei backbones were frozen and used solely as feature extractors.

The feature encodings produced by the Malinois and Sei backbones were pooled separately using custom attention-weighted networks to reduce dimensionality, and subsequently fused using a gated fusion mechanism. The resulting fused representation was mapped to MPRA activity using a single linear output layer. The model was trained for 50 epochs using the AdamW optimizer^98^ with a cosine learning rate scheduler.

To prevent data leakage due to sequence similarity across splits, data partitioning was performed at the virus strain level. Virus strains were randomly assigned to the training, validation, or test sets, ensuring that all tile sequences originating from a single strain were strictly contained within a single set. Consequently, the training set consisted of 49,316 tiles derived exclusively from double-stranded DNA viruses (**Table S7**). The validation and test sets comprised 8,653 and 7,059 tiles, respectively, and included both double-stranded DNA and retroviral sequences.

### LARM model architecture

To predict the fold-activation of HIV-1 LTR core promoter tile (HXB2:250-431) variants in response to TNFα treatment in Jurkat cells, we developed a separate model termed LARM (LTR Activation Response Model). The CREST model could not be applied to this task due to the structure of the experimental dataset. Specifically, the TNFα fold-activation dataset comprised 1,792 highly similar 180–200 bp sequences representing variants of the LTR core promoter region across HIV-1 isolates. This small and redundant dataset is insufficient for training a general sequence-to-activity deep learning model.

Therefore, we employed a simpler, two-step machine learning approach guided by prior biological knowledge. Based on existing literature, we expected the LTR core promoter to be primarily regulated by transcription factors (TFs) from the SP, RELA, ATF, ETS, and USF families. Consequently, the first step of LARM involves generating Sei predictions of the ChIP-seq tracks corresponding to these specific TFs. To obtain Sei predictions for a specific tile, the tile sequence is flanked with the MPRA plasmid sequence to achieve the input size required by Sei. In the second step, a Gradient Boosting Decision Tree (GBDT) algorithm is trained to predict TNFα fold-activation using these Sei-derived TF predictions as input features. We implemented the GBDT in Python using the open-source LightGBM package^99^. We randomly split the TNFα dataset into 1,254 training, 269 validation, and 269 test tiles (**Table S7**). We manually fine-tuned the

GBDT hyperparameters and used root mean square error (RMSE) as the objective function. The resulting model achieved Pearson correlation coefficient (PCC) values of approximately 0.9 on both the validation and test sets. Full implementation and training details are available on GitHub (https://github.com/osyafinkelberg/mpra-predictor).

### Longitudinal cohort evaluation of LTR activity

To study the longitudinal evolution of plasma HIV-1 LTR activity, we utilized data generated by Zanini et al.^65^ In this study, the authors performed whole-genome sequencing of HIV-1 populations from nine untreated PWH, with 6-12 longitudinal samples per PWH spanning 5-8 years of infection. For each sample, the HIV-1 genomes were amplified using PCR fragments spanning the entire genome, subject to further RNA sequencing with 250-300 nucleotide reads. As the starting point for our analysis, we used the provided .bam files (available at https://hiv.biozentrum.unibas.ch/) containing reads mapped to PWH-specific consensus HIV-1 genomes, which represent the reconstructed founder isolates.

We focused on several regions defined by HXB2 isolate coordinates, including the LTR core promoter (250–450 bp) and the internal CRE sites (7432–7632 bp and 7712–7912 bp). Each region was processed using the same pipeline, as described below.

The location of each region of interest within a PWH-specific consensus genome was identified using local Smith-Waterman alignment implemented in the Edlib Python package (https://pypi.org/project/edlib/). We considered reads from all PCR amplicons overlapping the region and used the Pysam Python library (https://github.com/pysam-developers/pysam) to extract unique sequence variants from reads completely covering the region. We refer to these unique variants as haplotypes, following the terminology of Zanini et al. During extraction, we allowed for variable haplotype lengths exceeding 200 bp. If a haplotype was shorter than 200 nucleotides due to indels, we padded the sequence with flanking read nucleotides to meet the 200 bp length requirement. To exclude artificial haplotype sequences arising from sequencing artifacts, we applied a frequency threshold, retaining only haplotypes with a frequency of at least 0.3% and supported by at least two different reads.

Phylogenetic analysis was performed independently for each PWH in three steps. First, we used MAFFT^96^ with the --auto strategy to realign all haplotypes that passed the frequency threshold. This alignment served as the input for maximum likelihood phylogeny reconstruction using FastTree^100^ under the Generalized Time-Reversible (GTR) nucleotide substitution model. Finally, IQ-TREE2^101^ was used to perform ancestral state reconstruction by applying the GTR substitution model and fixing the tree topology to the phylogeny predicted by FastTree.

Using the assembled sets of haplotype and ancestral sequences, we predicted basal MPRA activity in Jurkat cells and TNFα fold-activation using the CREST and LARM models, respectively (**Table S9**). For haplotype sequences longer than 200 bp, we generated tiling predictions for every 200 bp subsequence using a 1 bp step size, taking the maximum predicted activity value across all tiles. Substituting the maximum value with the average or median values did not qualitatively change the results. To quantitatively estimate the predicted activity change as a function of evolutionary time, we performed a root-to-tip regression for each PWH. Specifically, we calculated the distance from the root (measured in substitutions per site) for each observed haplotype (leaf node) on the PWH-specific phylogenetic trees. We then applied Ordinary Least Squares (OLS) linear regression to model the predicted activity against this tree depth. PWH with fewer than five observed haplotypes were excluded from this analysis. To rigorously assess the statistical significance of the observed evolutionary trends, we utilized a permutation test. The predicted activity values were randomly shuffled 100,000 times to break the phylogenetic association, allowing us to calculate an empirical P-value defined as the proportion of null slopes more extreme than the observed slope. Finally, to explain these changes in activity, we performed sequence-based motif annotation following the procedure described in the “Motif grammar analysis in HIV-1 sequences” section.

### Analysis of LTR activity across transmission pairs

To investigate the functional impact of HIV-1 LTR variation during vertical transmission, we analyzed data from Mehta et al.^62^ In this study, viral DNA was obtained from peripheral blood mononuclear cells (PBMCs) from six HIV-1–infected mother–infant pairs, cloned and sequenced, yielding 8–15 LTR variants per individual (total of 218 LTR sequences).

To characterize the genetic and phenotypic properties of sexually transmitted HIV-1 variants, we analyzed data from Claiborne et al.^63^ This study included six HIV-1 subtype C heterosexual transmission pairs from the Zambia-Emory HIV Research Project (ZEHRP) cohort in Lusaka, Zambia, with recipients additionally followed in the International AIDS Vaccine Initiative (IAVI) Protocol C early-infection cohort. Epidemiological linkage between partners was confirmed in the original study by phylogenetic analysis of HIV-1 gp41 sequences. From this dataset, 154 sequences were used in our downstream prediction analyses. We focused on the region defined by HXB2 isolate coordinates 250-450 bp, which include the LTR core promoter/enhancer. Ancestor sequence for child sequence was reconstructed and CREST was applied as described in the previous section of longitudinal cohort evaluation of LTR activity.

### Prediction of intragenic CREs using CREST

To obtain a comprehensive genome-wide annotation of CRE activity across diverse HIV-1 and HIV-2 isolates, we downloaded the 2022 curated filtered multiple sequence alignments of HIV-1 and HIV-2 complete genomes from the Los Alamos HIV Database (https://www.hiv.lanl.gov/content/sequence/NEWALIGN/align.html). We developed a custom script to iterate over the alignment using a 10 bp step size for HIV-1 and 1 bp step size for HIV-2. At each alignment position, we extracted a 200-nucleotide sequence from the ungapped genome of each individual isolate. For each extracted sequence, we predicted Jurkat MPRA activity using the CREST model and forward-strand CAGE-seq track expression (“FANTOM_CAGE_fwd”) using the Puffin model.^102^ Finally, we excluded isolates that lacked predicted activity at more than 40% of positions due to the presence of unknown nucleotides (N) in their genomic sequences. For visualization and reporting purposes, the resulting genome-wide activity maps were anchored to the coordinates of the HXB2 and SIV-Mac239 reference isolate genomes for HIV-1 and HIV-2, respectively. This anchoring was achieved by mapping the coordinates back to the aligned position of the reference sequence included in the multiple sequence alignment.

### Prediction of motifs contributing to activity within active regions

We generated a TF binding site (TFBS) annotation of HIV-1 CREs based on CREST hypothetical contribution scores, adapting a previously described procedure^27^ briefly outlined below. First, we calculated contribution scores for the CREST Jurkat MPRA predictions using a custom implementation of the Sampled Integrated Gradients (SIG) method.^26^ This was performed on a collection of 182,761 human Jurkat-specific DNase Hypersensitivity Sites (DHSs) obtained from ENCODE (ENCFF262WSM). We utilized this extensive set of CREs to maximize the recall of the TF-MoDISco-lite motif discovery algorithm.^103^ Next, we manually assigned TF motif families to the 33 distinct patterns identified by TF-MoDISco-lite. Finally, the resulting motif matrices were used to scan the contribution score profiles of the HIV-1 CREs (**Table S10**). This scanning was performed using the Continuous Jaccard Similarity (CJS) measure,^103^ consistent with our previous work.^27^

### Alphafold3 prediction of p65-p50-SP1 binding to HXB2:250-431

Protein-DNA multimeric complexes were predicted using AlphaFold3 multimer mode.^104^ Human NF-κB p65 (RelA; UniProt: Q04206 · TF65_HUMAN) and NF-κB p50 Rel homology domains (RHDs; NFKB1, UniProt: P19838 · NFKB1_HUMAN) were supplied as separate protein chains together with SP1 (UniProt: P08047 SP1_HUMAN), and paired with a double-stranded HIV-1 REJO LTR sequence “GAAGTTTGACAGCAGGCTAGCGTTTCATCACATGGCCCGAGAGAAGCATCCGGAGTACTTCAAGGACTGCTGACACCGAGCTTTCTTTGCTGACATCGAGCTTTCTACAAGGGACTTTCCGCTGGGACTTTCCGGGGAGGCGTGACCTGGGCGGGACTGGGGAGTGGCGAGCCCTCAGAAGCTGCATATAA” in a single multimer prediction run. The modeled complex consisted of two p65–p50 heterodimers and three SP1 molecules, with stoichiometry informed by the number of NF-κB and SP1 binding sites identified within the REJO LTR tile HXB2:250-431 sequence in saturation mutagenesis MPRA experiments, and intended to reflect potential multi-site occupancy. To minimize boundary effects and focus on the core regulatory sequence involving NF-κB and SP1 binding sites, the 331st to 431st basepairs of the REJO LTR tile HXB2:250-431 region was used. For each input configuration, five multimer models were generated and ranked using per-chain pLDDT, inter-chain predicted aligned error (PAE), and AlphaFold3’s interface confidence metrics (iPTM and chain-pair interface scores). The top-ranked model was selected for downstream interpretation. Structural visualization was performed using UCSF ChimeraX.^105^

### CASCADE microarray design

A custom microarray was designed to profile viral DNA elements (Agilent Technology Inc., AMADID 086310 and 086772, 4x180K format). Microarray probes contain a 24 nt (nucleotide) constant primer region, a 34 nt variable region, and a 5’- GC dinucleotide cap (**Table S13**). The microarray contained three classes of DNA probes: (1) tiled viral DNA elements (24,600 probes) (2) HIV-1 isolate sequences of target viral regulatory elements (9710 probes) ; (3) exhaustive mutagenesis of target viral regulatory elements (111,390 probes). Tiled viral DNA element probes capture 92 viral DNA elements covered by 26-nt-long DNA probe sequences tiled every 5-nt. Exhaustive mutagenesis probes were used to determine DNA binding motifs for 12 target regulatory elements (**Table S13**). As with the tiled probes, these regulatory elements are tiled every 5-nt using 26-nt-long probe sequences; however, for each 26-nt-long region we also included probes for all 3x26 single nucleotide variants. We also included 2610 random DNA probes to assess binding in relation to background. All unique probe sequences were included in two orientations on the microarray, and in 5-fold replicates each.

### Cell preparation for CASCADE

Jurkat cells were grown in RPMI media with 10% Fetal Bovine Serum (R&D Systems, Catalog # S12450H) and 1% Antibiotic-antimicotic (Thermofisher Scientific, Catalogue #15240062) up to a density of 1 million cells per mL prior to stimulation. Cells were stimulated with 10 ng/mL PMA (Millipore Sigma, Cat# P8139-1MG) and 1 µg/mL of anti-CD3 (BioLegend, Cat# 317302) in 100 mL for 1 hour. After 1 hour, they were processed for nuclear extraction.

### Nuclear extraction for CASCADE

Approximately 100 million Jurkat cells were harvested for each nuclear extraction replicate. Cells were collected in a falcon tube, placed on ice, and pelleted by centrifugation at 500 × g for 5 min at 4°C. Cell pellets were resuspended in 10 mL of 1X phosphate-buffered saline (PBS) with protease inhibitor and pelleted again at 500 × g for 5 min at 4°C and PBS wash was removed. To lyse the plasma membrane, cells were resuspended in 1.5 mL of Buffer A [10 mM HEPES, pH 7.9, 1.5 mM MgCl, 10 mM KCl, 0.1 mM protease inhibitor, phosphatase inhibitor (Santa Cruz Biotechnology, catalog # sc-45044), 0.5 mM dithiothreitol (DTT; Sigma–Aldrich, catalog # 4315)] and incubated for 10 min on ice. Then, Igepal detergent (final concentration of 0.1%) was added to the cell and Buffer A mixture and vortexed for 10 s. To separate the cytosolic fraction from the nuclei, the sample was centrifuged at 500 × g for 5 min at 4°C to pellet the nuclei. The pelleted nuclei were then resuspended in 100 μl Buffer C [20 mM HEPES, pH 7.9, 25% glycerol, 1.5 mM MgCl, 0.2 mM ethylenediaminetetraacetic acid (EDTA), 0.1 mM protease inhibitor, phosphatase inhibitor, 0.5 mM DTT and 420 mM NaCl] and then vortexed for 30 s. To extract the nuclear proteins (i.e. the nuclear extract), the nuclei were incubated in Buffer C for 1 hour while mixing at 4°C. To separate the nuclear extract from the nuclear debris, the mixture was centrifuged at 21 000 × g for 20 min at 4°C. The nuclear extract was collected in a separate microcentrifuge tube and flash frozen using liquid nitrogen. Nuclear extracts were stored at −80°C until used for CASCADE experiments.

### CASCADE experiment

All CASCADE protein-binding microarray (PBM) experiments were performed using the 4-chambered, 4x180K Agilent microarray platform (design details described below). DNA microarrays were double stranded as described.^106^ PBM experiments using cell extracts were performed following the protocols previously described.^107,108^ Briefly, the double-stranded microarray was pre-wetted in HBS + TX-100 (20 mM HEPES, 150 mM NaCl, 0.01% Triton X-100) for 5 min and then de-wetted in an HBS bath. Each of the microarray chambers was then incubated with 180 μL of nuclear extract binding mixture for 1 h in the dark. Nuclear extract binding mixture (per chamber): 400–600 μg of nuclear extract; 20 mM HEPES (pH 7.9); 100 mM NaCl; 1 mM DTT; 0.2 mg/mL BSA; 0.02% Triton X-100; 0.4 mg/mL salmon testes DNA (Sigma-Aldrich, Catalog #D7656). The microarray was then rinsed in an HBS bath containing 0.1% Tween-20 and subsequently de-wetted in an HBS bath. After the nuclear extract incubation, the microarray was incubated for 20 min in the dark with 20 μg/mL primary antibody for the TF or cofactor of interest diluted in 180 μL of 2% milk in HBS. The following primary antibodies were used to probe factors on the arrays: SP1 (Santa Cruz Biotechnology, Cat# sc-420), NCOA3 (Abcam, Cat# A300-347A), p300 (Abcam, Cat# ab14984), JMJD2A (Santa Cruz Biotechnology, sc-271210) and RBBP5 (Abcam, Cat# A300-109A). These antibodies were validated by western blot. After the primary antibody incubation, the array was first rinsed in an HBS bath containing 0.1% Tween-20 and then de-wetted in an HBS bath. Microarrays were then incubated for 20 min in the dark with 10 μg/mL of either Alexa488- or Alexa647-conjugated secondary antibody diluted in 180 μL of 2% milk in HBS. The following fluorescently labeled secondary antibodies were used in our CASCADE experiments, and were species matched to the primary antibodies described above: Goat anti-rabbit IgG (H + L) Highly Cross-absorbed Secondary Antibody, Alexa Fluor 647 (ThermoFisher, Cat #A32733); Goat anti-mouse IgG (H + L) Highly Cross-absorbed Secondary Antibody, Alexa Fluor 488 (ThermoFisher, Cat # A11029). Excess antibody was removed by washing the array twice for 3 min in 0.05% Tween-20 in HBS and once for 2 min in HBS in Coplin jars as described above. After the washes, the microarray was de-wetted in an HBS bath. Microarrays were scanned with a GenePix 4400 A scanner and fluorescence was quantified using GenePix Pro 7.2. Exported fluorescence data were normalized with MicroArray LINEar Regression^109^.

## Supplementary Figures

**Supplementary Figure 1:**
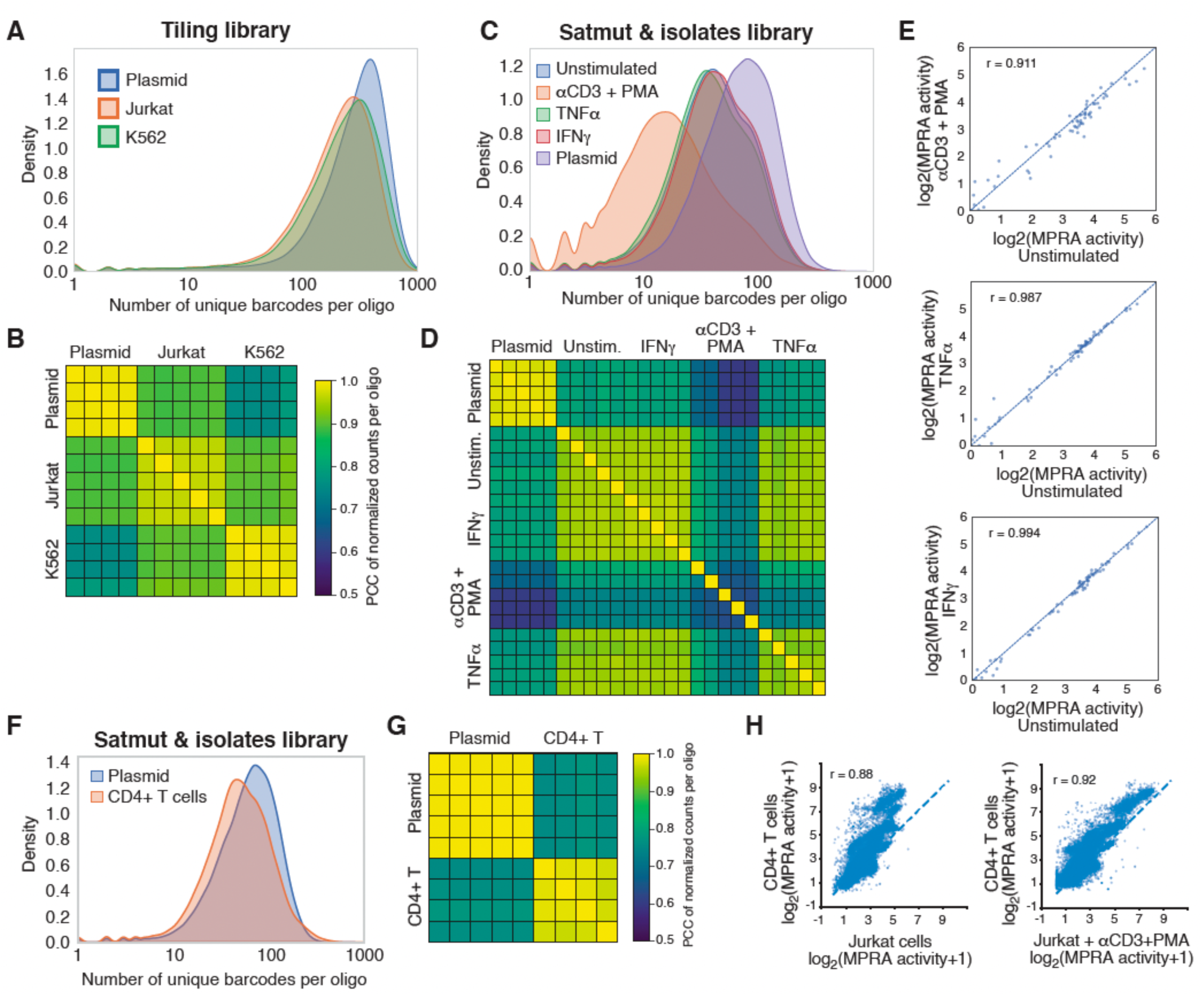
MPRA quality controls. (A, C) Distribution of number of barcodes per oligonucleotide in the plasmid pool and in the RNA fractions from different cell lines or conditions for the tiling library (A) or the isolate/saturation mutagenesis library (C). (B, D) Correlation between replicates and across samples for the tiling (B), and isolate/saturation mutagenesis (D) libraries. The Pearson correlation coefficients (PCC) between normalized read counts per oligonucleotide are shown. (E) Correlation in MPRA activity for human sequences known to be active by MPRAs between unstimulated Jurkat cells and Jurkat cells stimulated with αCD3+PMA, TNFα, or IFNγ. The diagonal shows the identity line. Correlation determined using Pearson correlation coefficient (r). (F) Distribution of number of barcodes per oligonucleotide in the plasmid pool and in the RNA fraction from CD4+ T cells from four donors for the isolate/saturation mutagenesis MPRA library. (G) Correlation between replicates and across donors for the isolate/saturation mutagenesis library. The Pearson correlation coefficients (PCC) between normalized read counts per oligonucleotide are shown. (H) Correlation between the MPRA activity determined in CD4+ T cells and the activity determined in unstimulated or αCD3+PMA-stimulated Jurkat cells. The Pearson correlation coefficient is indicated.

**Supplementary Figure 2:**
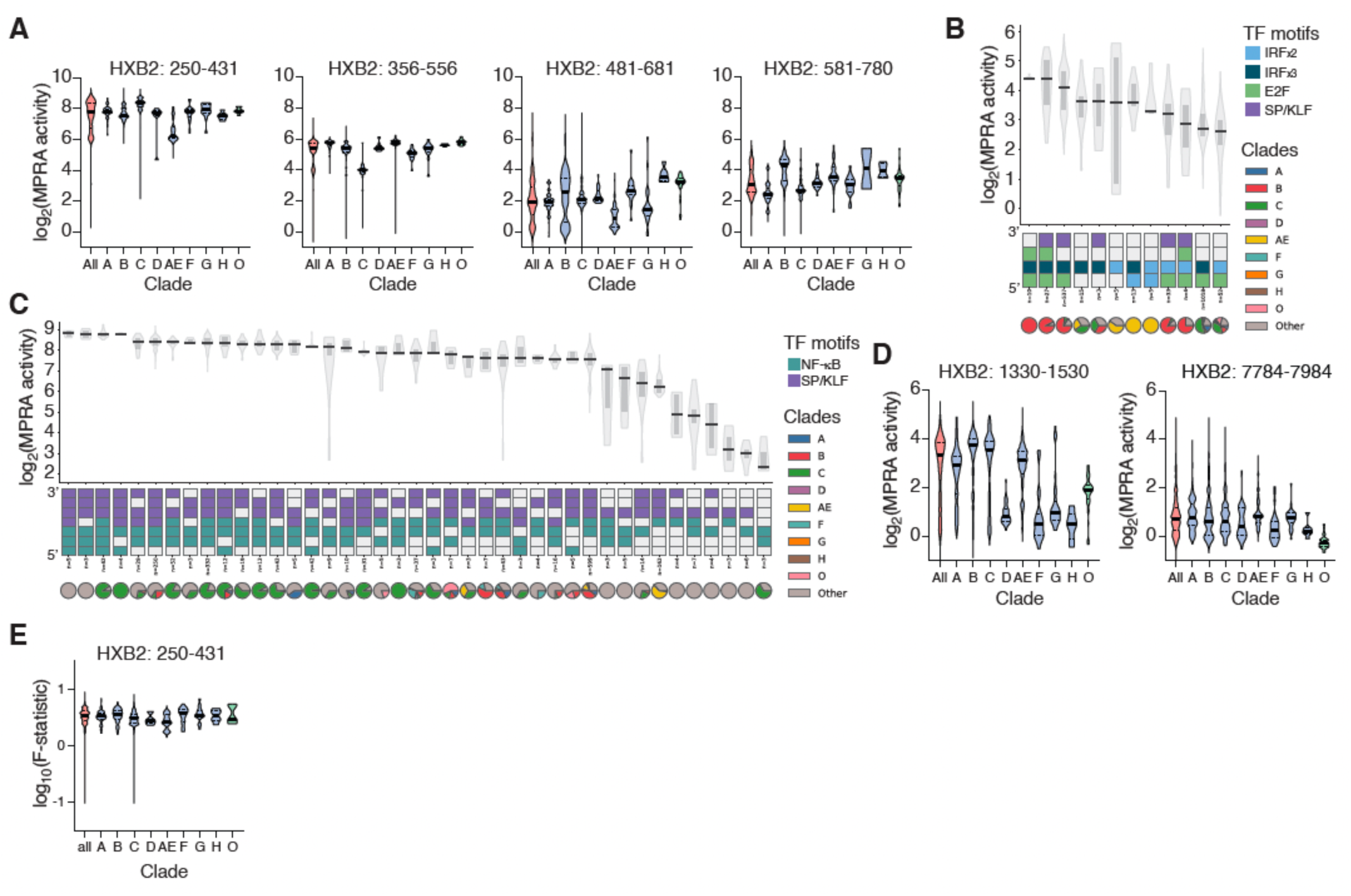
Variation in HIV-1 transcriptional activity in primary CD4+ T cells. (A) Violin plots showing the distribution of activity in CD4+ T cells across 5,569 isolates for different regions of the HIV-1 LTR across clades. The thick black line indicates the median, and the dotted lines indicate the first and third quartiles. (B-C) Activity distribution in CD4+ T cells cells across HIV-1 isolates with different TF configurations in tiles HXB2:250-431 (B) and HXB2:581-780 (C). The bottom boxes represent the TF configurations based on aligned TF positions. The distribution of isolates from different clades across TF configurations is shown as pie charts. Violin plots indicate the distribution of activity. (D) Violin plots showing the distribution of activity across isolates for intragenic CREs from HIV-1 across clades for tiles HXB2:1330-1530 and HXB2:7784-7984. The thick black line indicates the median, and the dotted lines indicate the first and third quartiles. (E) Violin plots of F-statistic showing the variability in activity for each isolate for tile HXB2:250-431 across four donors in CD4+ T cells. The thick black line indicates the median, and the dotted lines indicate the first and third quartiles.

**Supplementary Figure 3:**
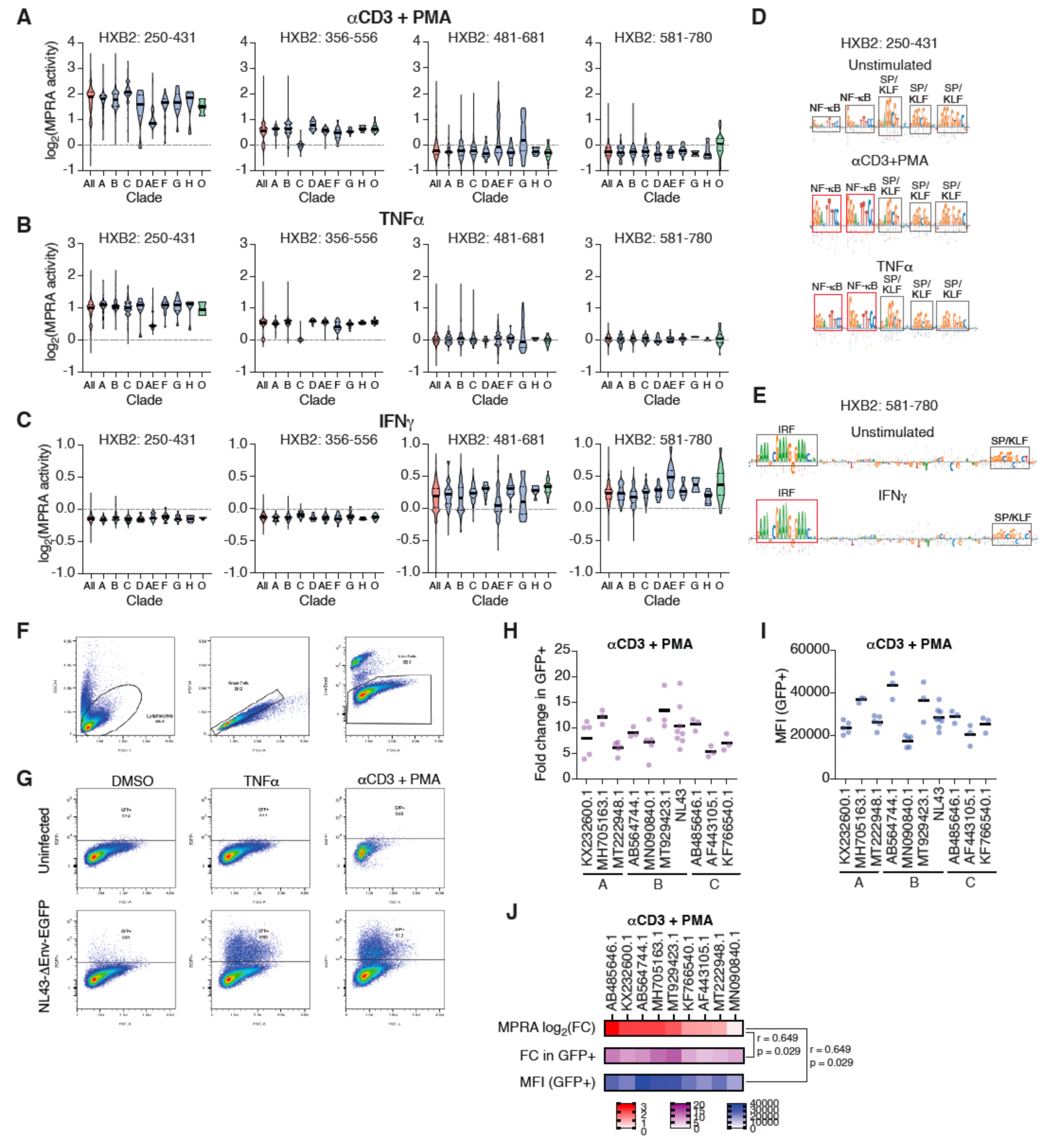
HIV-1 LTR activation across clades and isolates. (A-C) Violin plots showing the distribution of MPRA fold activation across HIV-1 clades and isolates in Jurkat cells stimulated with αCD3+PMA (A), TNFα (B), or IFNγ (C). The thick black line indicates the median, and the dotted lines indicate the first and third quartiles. (D) Comparison between contribution scores derived from MPRA saturation mutagenesis in unstimulated, αCD3+PMA, and TNFα treated Jurkat cells for tile HXB2:250-431. TF motifs are shown and the activated NF-κB motifs are outlined in red. (E) Comparison between contribution scores derived from MPRA saturation mutagenesis in unstimulated and IFNγ treated Jurkat cells for tile HXB2:581-780. TF motifs are shown and the activated IRF motif is outlined in red. (F) Cell gating used to evaluate transcription from variant LTR-HIV-ΔEnv-EGFP constructs. Sample corresponding to uninfected cells. (G) EGFP levels in cells uninfected or infected with NL43-ΔEnv-EGFP, stimulated with DMSO, TNFα, or αCD3+PMA. (H-I) Fold change in GFP+ cells relative to DMSO control (H) or geometric mean fluorescence intensity of GFP+ cells (I) in Jurkat cells infected with chimeric proviruses carrying the U3 region of the indicated isolates from clades A, B, and C, and stimulated with αCD3+PMA for 2 days. (J) Comparison between fold activation of MPRA data from tile HXB2:250-431 in Jurkat cells activated with αCD3+PMA, and provirus reactivation by αCD3+PMA measured as fold change in GFP+ cells relative to DMSO control or geometric mean fluorescence intensity of GFP+ cells. Pearson correlation coefficients and one-tailed p-values are indicated for each comparison to MPRA data.

**Supplementary Figure 4:**
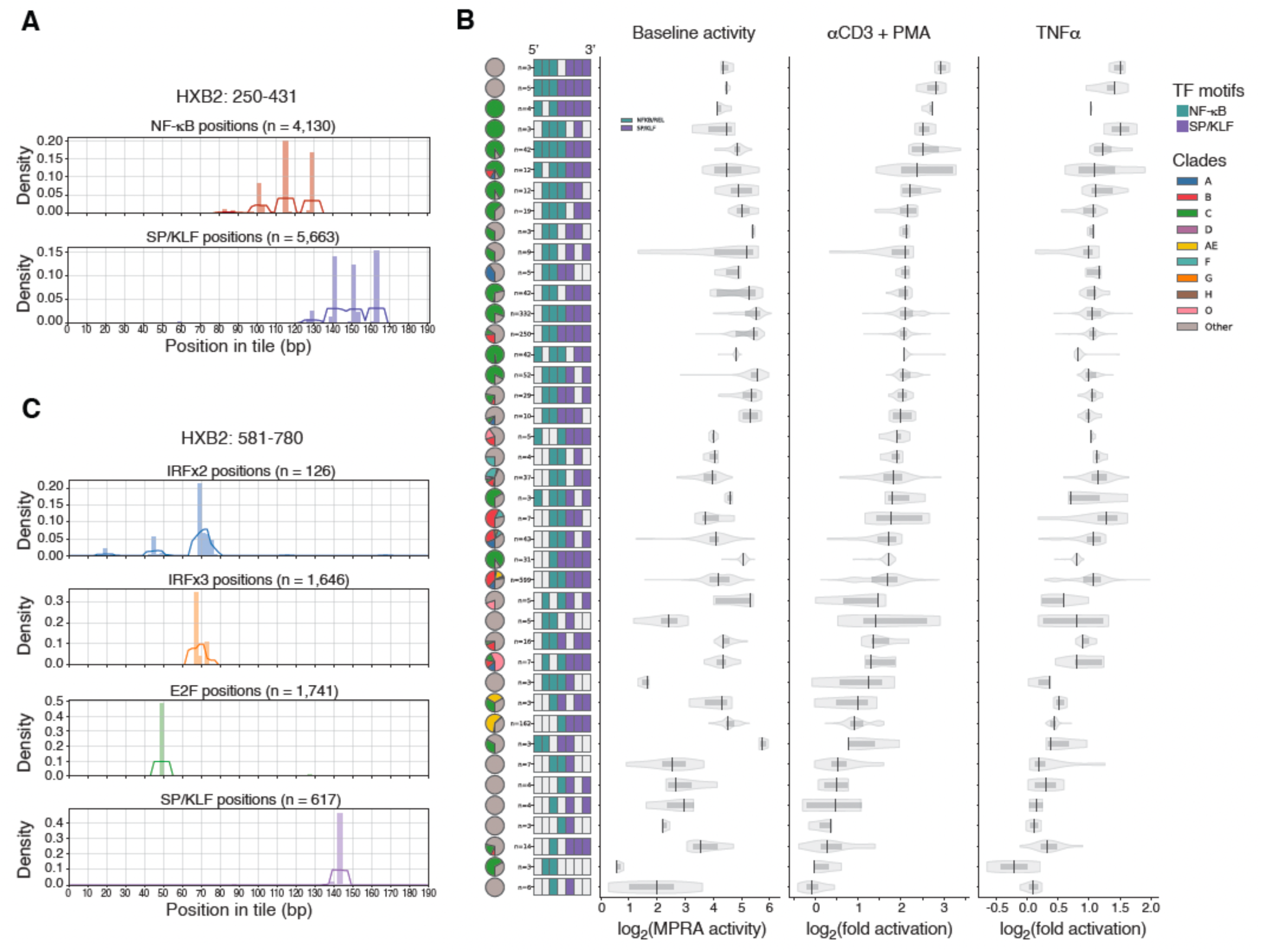
Influence of TF numbers and configuration on HIV-1 LTR activity. (A, C) Distribution of TF motifs across isolates aligned HIV-1 LTR sequences from tiles HXB2:250-431 (A) and HXB2:581-780 (C). (B) Activity distribution across HIV-1 isolates with different TF configurations in tile HXB2:250-431. The left boxes represent the TF configurations based on aligned TF positions. The distribution of isolates from different clades across TF configurations is shown as pie charts. Violin plots indicate the distributions of activity in unstimulated Jurkat cells (baseline), and cells stimulated with αCD3+PMA or TNFα.

**Supplementary Figure 5:**
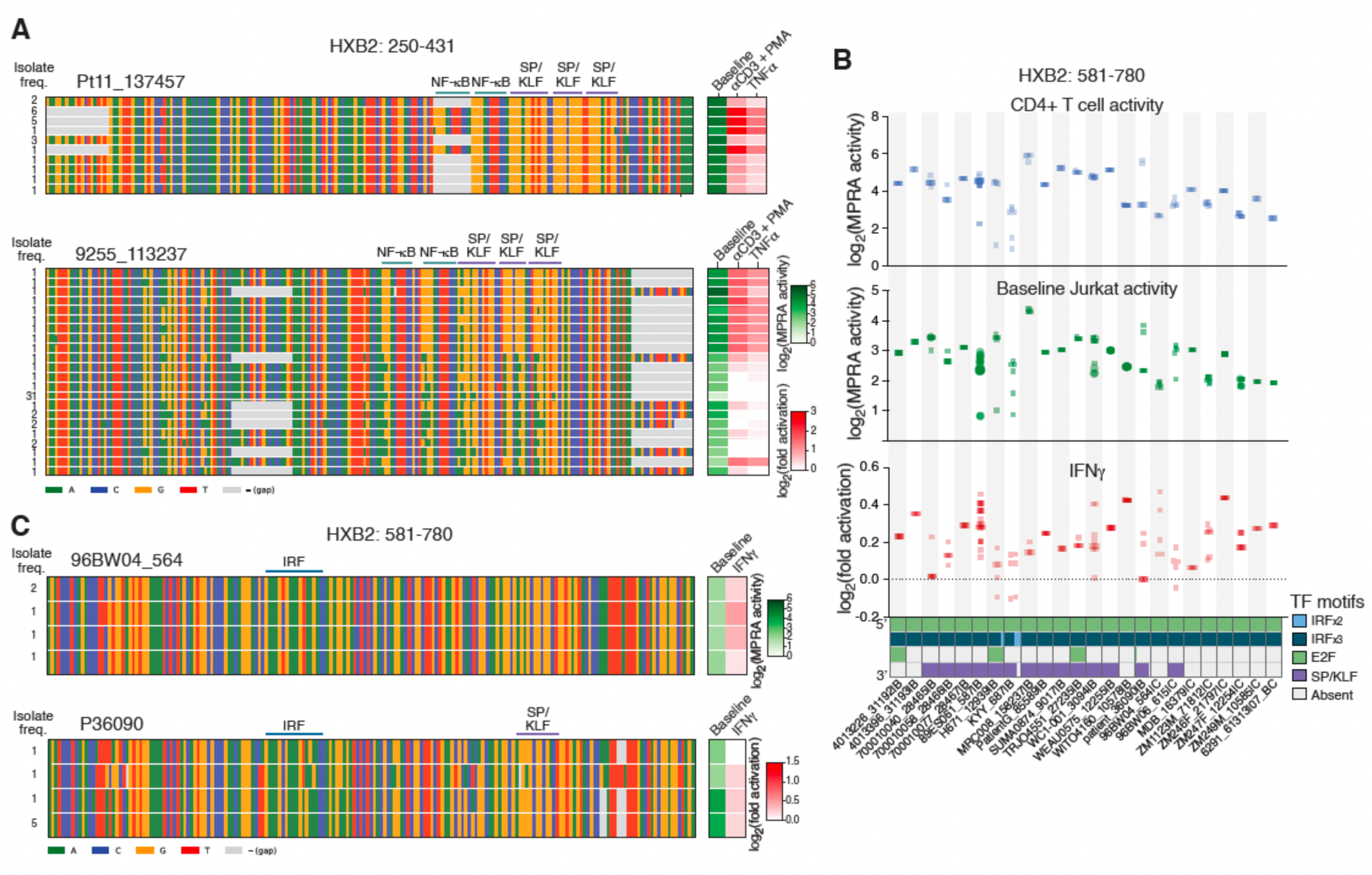
Variation in HIV-1 LTR activity and regulation within PWH. (A, C) Examples of activity and sequence variation for isolates from PWH corresponding to tiles HXB2:250-431 (A) or HXB2:581-780 (C). The left heatmaps show the sequence alignments for isolates of a PWH. The frequency of isolates with a specific sequence is shown in the left. TF binding sites are indicated on top. The right heatmap indicates the baseline activity (green) and fold activation by αCD3+PMA, TNFα, or IFNγ (red). (B) Distribution of baseline activity and fold activation by IFNγ for PWH with at least 5 different isolates in tile HXB2:581-780. PWH are ordered by HIV-1 clade. The bottom boxes represent the TF configurations based on aligned TF positions across all isolates from the corresponding PWH.

**Supplementary Figure 6:**
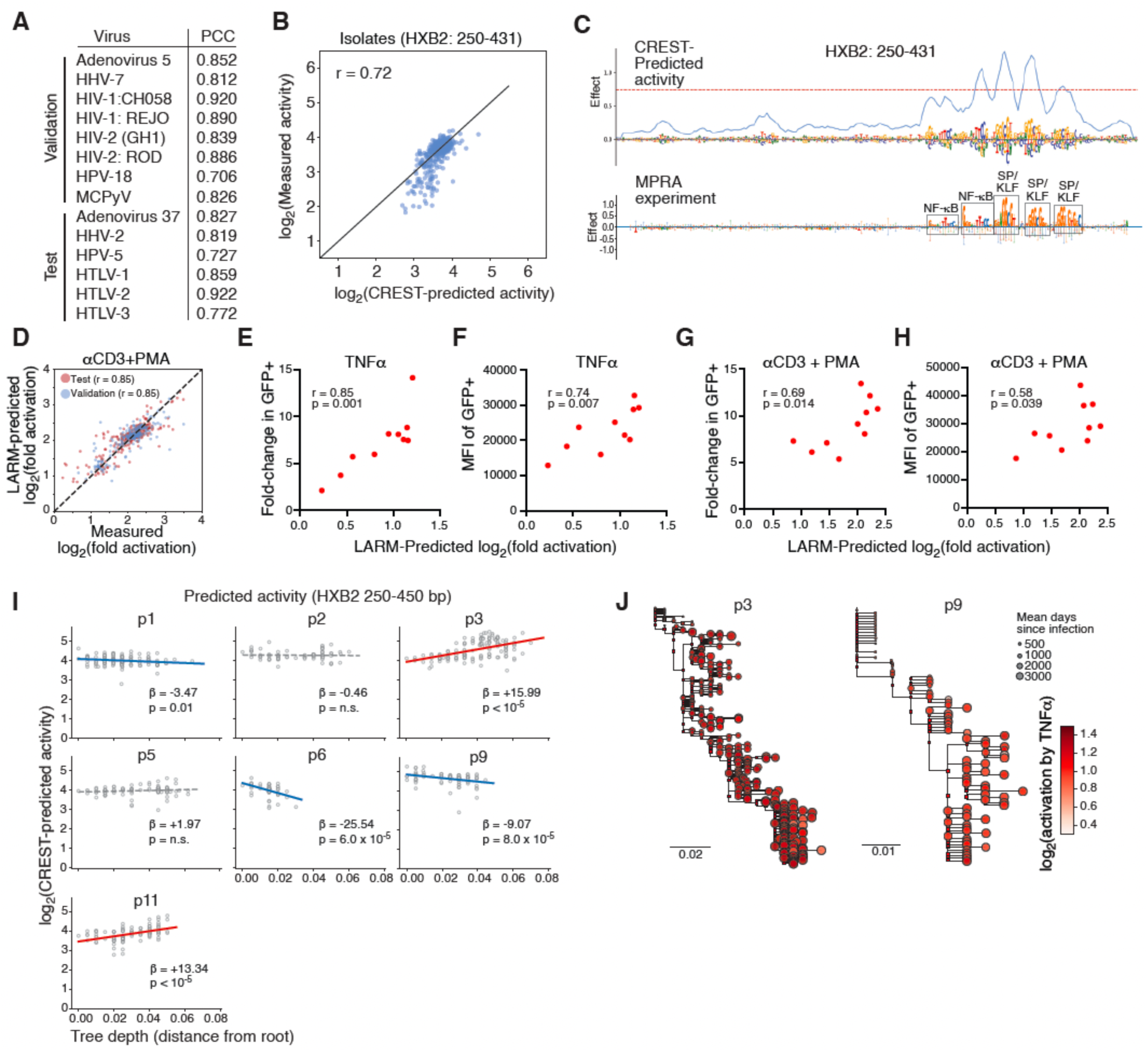
Training and validation of CREST and LARM models. (A) Tiles from the indicated viral genomes used as part of the validation and test sets. Pearson correlation coefficient (PCC) values between measured and predicted activities are shown for each virus. (B) Correlation between measured MPRA activity and baseline activity predicted using CREST for tile HXB2:250-431 across different isolates. The diagonal indicates the identity line. r = Pearson correlation coefficient. (C) Contribution scores for saturation mutagenesis based on predicted activity using CREST (top) compared to experimental MPRA saturation mutagenesis in Jurkat cells (bottom) for tile HXB2:250-431. TF motifs are outlined. (D) Pearson correlation for tile HXB2:250-431across isolates between fold activation induced by αCD3+PMA in Jurkat cells measured by MPRAs and the corresponding values predicted by LARM for the validation and test sets. (E-H) Correlation between fold activation by TNFα (E-F) or αCD3+PMA (G-H) predicted using LARM for tile HXB2:250-431 and measured fold change in GFP+ cells (E, G) or geometric mean fluorescence intensity of GFP+ cells (F, H) for seven HIV-1 isolates tested in the context of integrated proviruses. Pearson correlation coefficient (r) and significance (p) are indicated. (I) Predicted LTR haplotype activity regressed against tip-to-root distance in a maximum-likelihood phylogeny reconstructed using FastTree for 7 PWH.^65^ The p-value of the regression slope was calculated using a permutation test with 100,000 permutations. (J) Fold activation by TNFα predicted using LARM for tile HXB2:250-431 of two PWH. Isolates are shown within their respective evolutionary trees. Node color represents the activity and node size indicates the average day post infection for the corresponding haplotype. Square nodes correspond to predicted ancestral sequences.

**Supplementary Figure 7:**
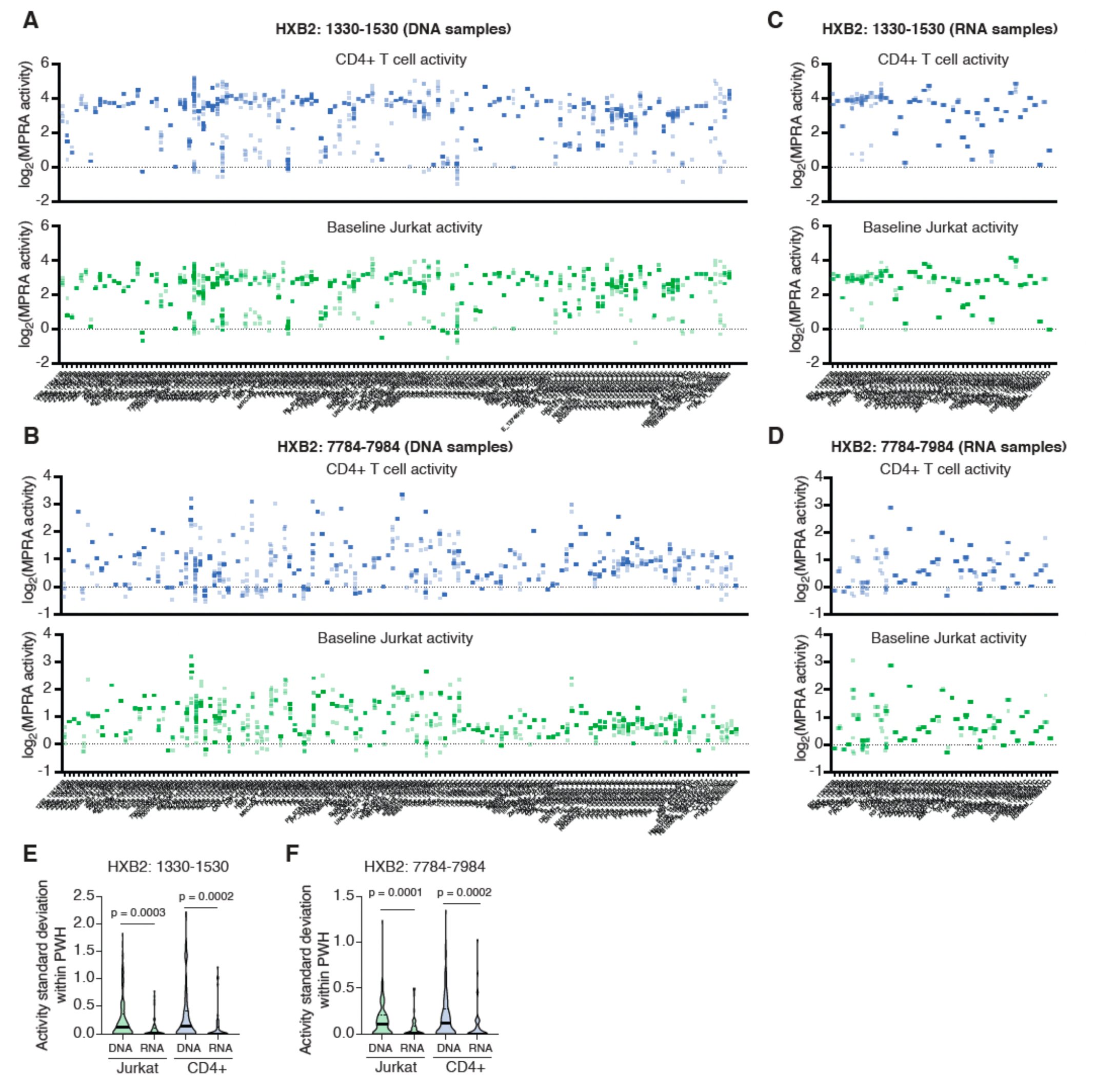
Variation in activity of HIV-1 intragenic CREs within PWH. (A-D) Distribution of MPRA activity in Jurkat cells and CD4+ T cells in PWH with at least 5 isolates for tiles (HXB2:1330-1530 (A, C) and HXB2:7784-7984 (B, D). (A-B) show sequences obtained from proviral DNA, whereas (C-D) show sequences obtained from viral RNA. (E-F) Violin plots showing the distribution of MPRA activity variation, measured as the standard deviation in activity for isolates within individuals, for DNA- and RNA-derived isolates, corresponding to tiles HXB2:1330-1530 (E) and HXB2:7784-7984 (F). The thick black line indicates the median, and the dotted lines indicate the first and third quartiles.

## Notes

### Competing Interest Statement

The authors have declared no competing interest.

